# Epigenetic age is accelerated in schizophrenia with age- and sex-specific effects and associated with polygenic disease risk

**DOI:** 10.1101/727859

**Authors:** Anil P.S. Ori, Loes M. Olde Loohuis, Jerry Guintivano, Eilis Hannon, Emma Dempster, David St. Clair, Nick J Bass, Andrew McQuillin, Jonathan Mill, Patrick F Sullivan, Rene S. Kahn, Steve Horvath, Roel A. Ophoff

**Affiliations:** University of California Los Angeles, Center for Neurobehavioral Genetics, Semel Institute for Neuroscience and Human Behavior, Los Angeles, CA, USA; Department of Genetics, University Medical Center Groningen, The Netherlands; Department of Psychiatry, University Medical Center Groningen, The Netherlands; University of North Carolina, Department of Genetics, Chapel Hill, NC, US; University of Exeter, University of Exeter Medical School, Exeter, UK; University of Aberdeen, Institute of Medical Sciences, Aberdeen, Scotland, UK; University College London, Division of Psychiatry, UK; King’s College London, London, UK; Karolinska Institutet, Department of Medical Epidemiology and Biostatistics, Stockholm, Sweden; Icahn School of Medicine at Mount Sinai, Department of Psychiatry, New York, NY, USA; University of California Los Angeles, Department of Biostatistics, Fielding School of Public Health, Los Angeles, CA, USA; University of California Los Angeles, Department of Human Genetics, David Geffen School of Medicine, Los Angeles, CA, USA; Erasmus University Medical Center, Department of Psychiatry, Rotterdam, The Netherlands

**Keywords:** schizophrenia, DNA methylation, aging, epigenetic clocks, biological aging, accelerated aging, polygenic risk, mortality risk

## Abstract

**Background:** The study of biological age acceleration may help identify at-risk individuals and contribute to reduce the rising global burden of age-related diseases. Using DNA methylation (DNAm) clocks, we investigated biological aging in schizophrenia (SCZ), a severe mental illness that is associated with an increased prevalence of age-related disabilities and morbidities. In a multi-cohort whole blood sample consisting of 1,090 SCZ cases and 1,206 controls, we investigated differential aging using three DNAm clocks (i.e. Hannum, Horvath, Levine). These clocks are highly predictive of chronological age and are known to capture different processes of biological aging.

**Results:** We found that blood-based DNAm aging is significantly altered in SCZ with age- and sex-specific effects that differ between clocks and map to distinct chronological age windows. Most notably, differential phenotypic age (Levine clock) was most pronounced in female SCZ patients in later adulthood compared to matched controls. Female patients with high SCZ polygenic risk scores (PRS) present the highest age acceleration in this age group with +4.30 years (CI: 2.40-6.20, P=1.3E-05). Phenotypic age and SCZ PRS contribute additively to the illness and together explain up to 22.4% of the variance in disease status in this study. This suggests that combining genetic and epigenetic predictors may improve predictions of disease outcomes.

**Conclusions:** Since increased phenotypic age is associated with increased risk of all-cause mortality, our findings indicate that specific and identifiable patient groups are at increased mortality risk as measured by the Levine clock. These results provide new biological insights into the aging landscape of SCZ with age- and sex-specific effects and warrant further investigations into the potential of DNAm clocks as clinical biomarkers that may help with disease management in schizophrenia.

## Introduction

As the population continues to age, reducing the burden of age-related disability and morbidity is timely and important, particularly for mental illnesses [1,2]. Ranked as one of the most disabling illnesses globally[3], schizophrenia (SCZ) has significant impact on patients, families, and society. SCZ is associated with a two- to threefold increased risk of mortality[4–6] and a 15 year reduction in life expectancy compared to the general population[7,8]. Despite elevated rates of suicide and other unnatural causes of death, most morbidity in SCZ is attributed to age-related diseases, such as cardiovascular and respiratory diseases and diabetes mellitus[5,9,10]. Processes of biological aging may therefore be accelerated in patients diagnosed with SCZ, either through an increased prevalence of age-related conditions or as a more integrated part of the illness [11]. Quantification of biological aging can help with identification of at-risk individuals or even prevention of age-related diseases [12,13]. While different aging biomarkers have been studied in SCZ, no clear demonstration of altered biological age has been shown [14]. The recent development of DNA methylation (DNAm) age predictors however offers new opportunities to study the phenomenon of aging in SCZ.

DNAm age predictors, or “epigenetic clocks’’, are biomarkers of ageing that generate a highly accurate estimate of chronological age, known as DNAm age [15–17]. The difference (Δage) between predicted DNAm and chronological age is associated with a wide-range of health and disease outcomes, including all-cause mortality [18–21], socioeconomic adversity and smoking[22], metabolic outcomes, such as body mass index (BMI) and obesity [23,24], and brain-related phenotypes, such as Parkinson’s disease, posttraumatic stress disorder, insomnia, major depressive disorder, and bipolar disorder [25–29]. As epigenetic signatures can be modifiable [30], DNAm-based predictors may have significant clinical utility. Studies of DNAm aging so far found limited to no evidence for altered biological age in either brain or blood in SCZ [31–34]. These studies, however, (i) consisted of small sample sizes and thus limiting the ability to detect a biological signal, (ii) used a single DNAm clock that may have not been most informative for aging studies of mental illnesses, and (iii) did not consider aging differences across the lifespan of patients. As morbidities in the SCZ population differ between older and younger individuals, and females and males [5], analyses of both age- and sex-specific effects is warranted and could identify differential aging patterns, nevertheless.

To investigate DNAm aging in SCZ, we used three independent DNAm age estimators; the Hannum [16], Horvath [15], and Levine clock [17]. Each clock is designed using different training features and captures distinct characteristics of aging [35]; (i) the Hannum age predictor was trained on whole blood adult samples, (ii) the Horvath predictor was trained across 30 tissues and cell types across developmental stages, and (iii) the Levine combines DNAm from adult blood samples with clinical blood-based measures. As the Levine estimator is trained on chronological age and nine clinical markers, its output is referred to as DNAm PhenoAge or “phenotypic age”. The Hannum estimator is said to capture measures of cell extrinsic aging in blood, whereas the Horvath clock measures more cell intrinsic aging as it was trained across multiple tissues and therefore is less dependent on cell type composition. All three clocks, in different but complementary ways, capture the pace of biological aging that is associated with various age-related conditions and diseases, including all-cause mortality [19,35].

DNAm clocks were implemented across four European case-control cohorts, representing a sample of almost twice the size of the largest SCZ DNAm age study conducted so far. Analyses are performed across the full sample and stratified by age and sex. We then integrated DNAm age with age of onset, duration of illness, and SCZ polygenic risk. DNAm smoking scores and blood cell type proportions were used to gain further insights into differential aging patterns. This study overall reports an in-depth investigation of the DNAm aging landscape in schizophrenia.

## Results

Figure 1 shows a schematic overview of the study design and analysis framework used to investigate DNAm aging in SCZ. After data pre-processing and quality control, 1,090 SCZ cases and 1,206 controls (2,296 subjects of 2,707 initial samples) were included in our analysis. The overall sample has a mean age of 40.3 years (SD=14.4) and consists of 34.5% women (Table S1 and Figure S1).

**Figure 1.**
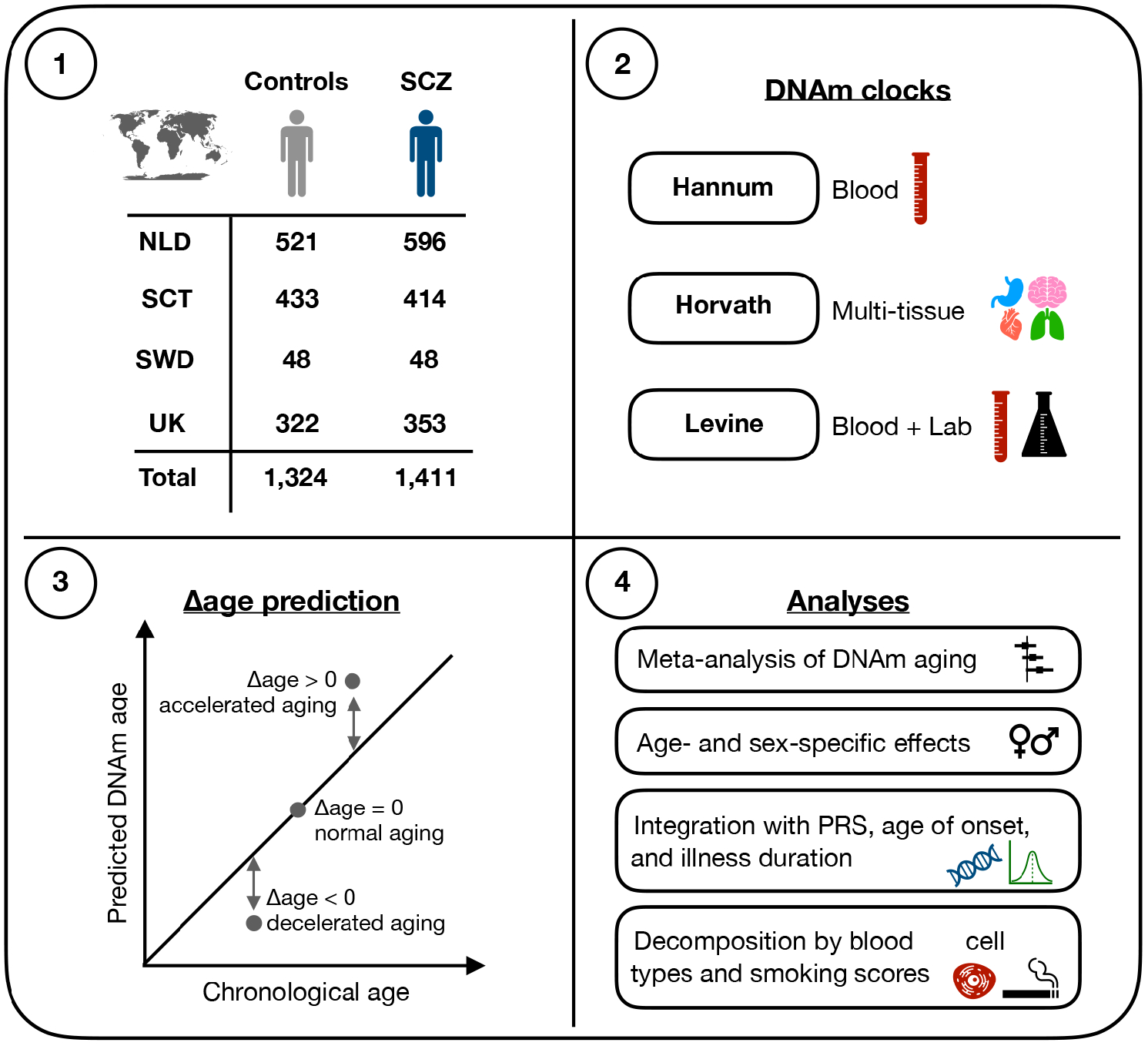
Overview of study design and analysis framework. DNA methylation (DNAm) data was available for a total of 2,735 samples across four European cohorts. See Table S2 for more details on samples. DNAm age estimates were generated using three DNAm clocks, each designed to capture different features of aging (box 2). To investigate differences in aging between cases and controls, Δage was computed (box 3) and analyzed according to the step-wise framework shown in box 4. SCZ = schizophrenia, NLD=Netherlands, SCT=Scotland, SWD=Sweden, UK=United Kingdom, PRS=polygenic risk scores.

Across cohorts, all three clocks produce a high correlation with chronological age (Pearson’s r = 0.92-0.94; Figure 2A and S2). Using duplicates in the Dutch cohort, we assessed consistency between pairs of technical replicates, i.e. samples for which blood was collected at the same time but DNA processed at different times and DNAm data obtained on different arrays. Comparing Δage estimates between these pairs, we find a significant correlation for each clock (Figure S3); Hannum (rho = 0.79, n = 10), Horvath (rho = 0.53, n=118), Levine (rho = 0.67, n=118). Δage directionality (i.e. age deceleration or acceleration) is concordant in 90%, 73%, and 86% of pairs for Hannum, Horvath, and Levine, respectively, highlighting that the obtained estimates of DNAm age are reproducible for all three clocks. Comparing Δage estimates between clocks using all samples, we find a moderate concordance (Pearson’s r = 0.39-0.43; Figure S4), demonstrating that a significant proportion of the variation in Δage is clock-specific. As these three estimators were trained on different features of biological aging, investigating them in conjunction may thus yield broader insights into differential aging.

**Figure 2.**
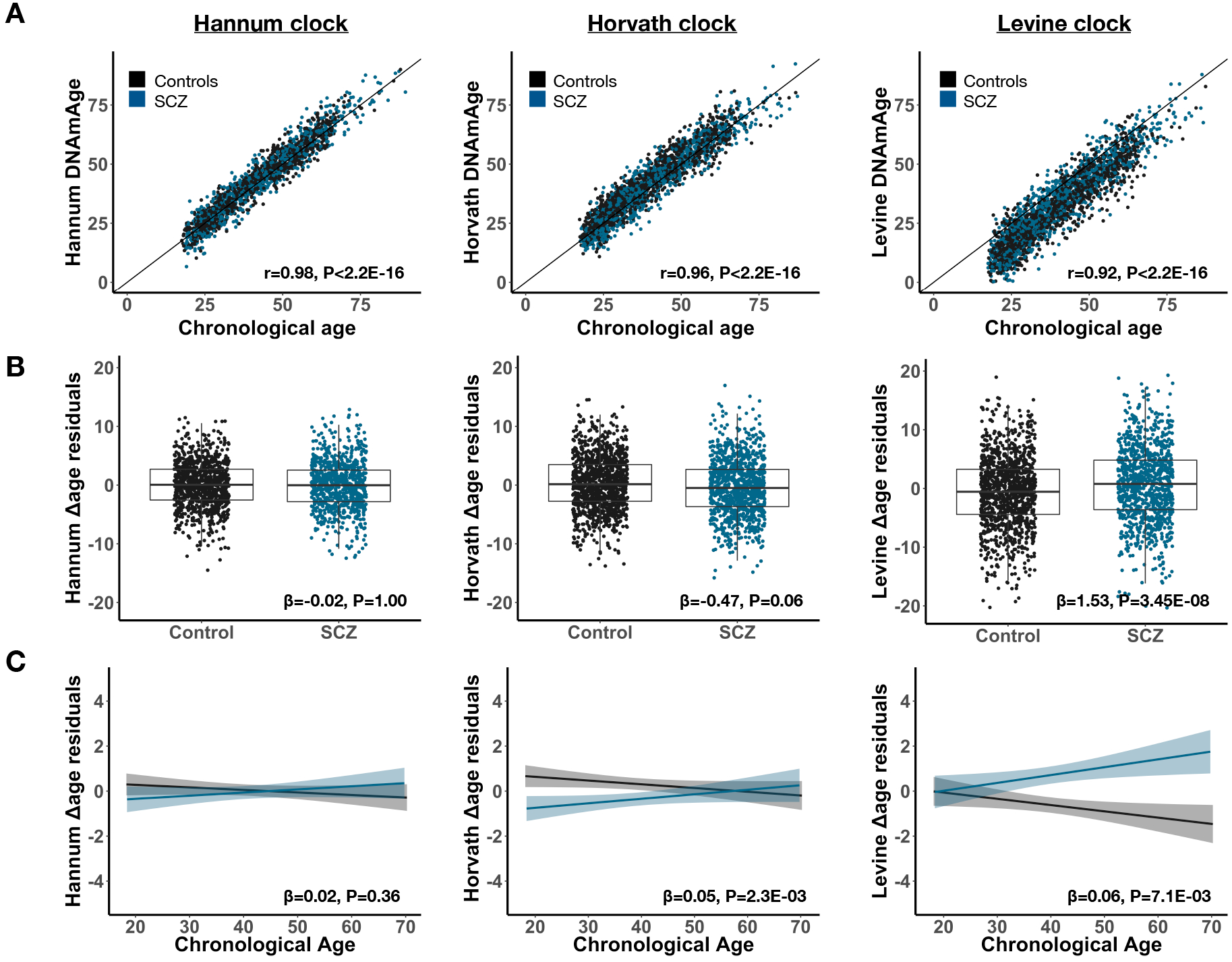
DNA methylation aging is altered in schizophrenia and conditional on chronological age. Presented are results visualizing DNAm aging in SCZ for each clock; Hannum (left), Horvath (middle), Levine (right). Cases are shown in blue and controls in black. (A) The correlation between DNAm age and chronological age. The Pearson’s correlation estimate and corresponding p-value are shown in the bottom corner. (B) Boxplots of Δage between cases and controls with the meta-analytic effect size and p-value across cohorts shown. β represents the mean change in Δage in cases compared to controls. (C) Δage is visualized across chronological age with a regression line fitted separately for cases and controls and the meta-analytic interaction effect and p-value shown. β represents the change in Δage in cases per year of chronological age compared to controls. P-values are adjusted for multiple testing across clocks (n=3).

### DNA methylation age is altered in an age-dependent manner

Across the full sample, patients with SCZ are on average 1.53 years older in phenotypic Δage (Levine clock) compared to controls (Pmeta= 3.45E-08) (Figure 2B). The intrinsic cellular age (Horvath) predictor revealed an opposite pattern, with SCZ cases appearing 0.47 years younger compared to controls (Pmeta= 0.06). No differences were observed between cases and controls when applying the blood-based Hannum DNAm age predictor. Within the analysis of each clock, we observed no evidence of heterogeneity between the four cohorts (Phet > 0.05, Table S5).

Modelling the interaction effect between disease status and chronological age on Δage reveals a differential rate of aging between cases and controls (Figure 2C). That is, the slope of Δage across chronological age is 0.05- and 0.06-years steeper in cases compared to controls for the Horvath (Pmeta=2.3E-03) and Levine clocks (Pmeta=7.1E-03), respectively, with no evidence of heterogeneity between cohorts (Figure S5 and Table S6). As no significant effects were observed for the Hannum Δage, we decided to focus our downstream analysis on the phenotypic (Levine) age and intrinsic cellular (Horvath) age only. To further disentangle the relationship between Δage in SCZ conditional on chronological age, we estimated differential aging by 10-year intervals, with years 18 and 19 included in the first age group. We observe significant DNAm age deceleration in early adulthood (18-30 years) with patients estimated at −1.23 years younger (Pmeta=3.9E-03) in intrinsic cellular age with no significant difference at later ages (Figure 3A). In phenotypic age, SCZ patients displayed significant DNAm age acceleration from 30 years and older (Figure 3B), with the most pronounced age acceleration between 50-60 years (2.29 years, Pmeta=9.0E-03). We again find no evidence of heterogeneity within age groups between cohorts (Figure S6 and Table S7-8).

**Figure 3.**
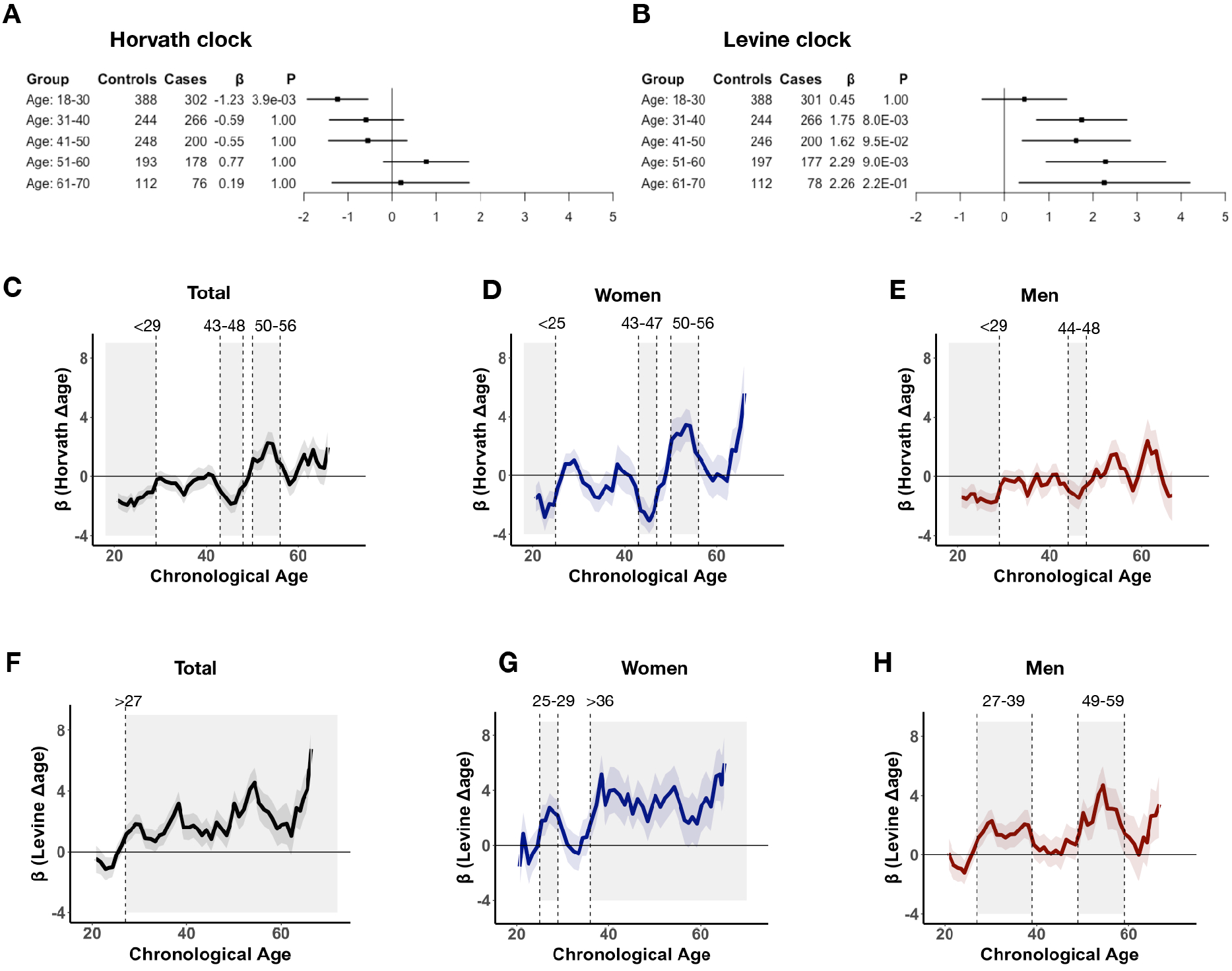
Differential DNAm aging in schizophrenia maps to specific age windows between sexes. (A-B) Shown are Δage differences between cases and controls across age groups for the Horvath (A) and Levine clock (B). For each age group, number of cases and controls, and meta-analytic effect size (β) and p-value (P) are presented. P-values are corrected for multiple testing (2 clocks x 5 groups = 10 tests). See Table S5 for more details on results and corresponding statistics. (C-H) Sliding age- windows, using 5-year bins with steps of 1-year, were used to estimate differential aging (β) at finer resolution across the range of chronological age. Significant shifts in Δage between cases and controls, defined by the standard error of β deviating from zero for at least 3 steps, are highlighted by the shaded areas on the graph with the dotted vertical lines indicating the respective ages of the intervals. Identified age intervals for the Horvath and Levine clock are shown in C-E and F-H, respectively. Results for women (middle) and men (right) are presented in blue and red, respectively. The effects in the total sample are displayed in black (left).

### Age- and sex-specific effects contribute to DNAm aging

To quantify the overall contribution of age- and also sex-specific effects, we estimated the gain in variance explained of Δage by adding the interaction terms of age and sex with disease status to a baseline model and assessed the gain in model performance. For both measures of aging, inclusion of interaction terms presented a significantly better fit, with the three-way interaction model (i.e. disease status, age and sex) explaining the most variance in Δage (Table 1 and S9). We observe a larger gain in model fit for the three-way interaction for phenotypic aging (P=0.01) than for intrinsic cellular aging (P=0.24), suggesting that sex-specific effects are more pronounced for Levine Δage.

**Table 1.**
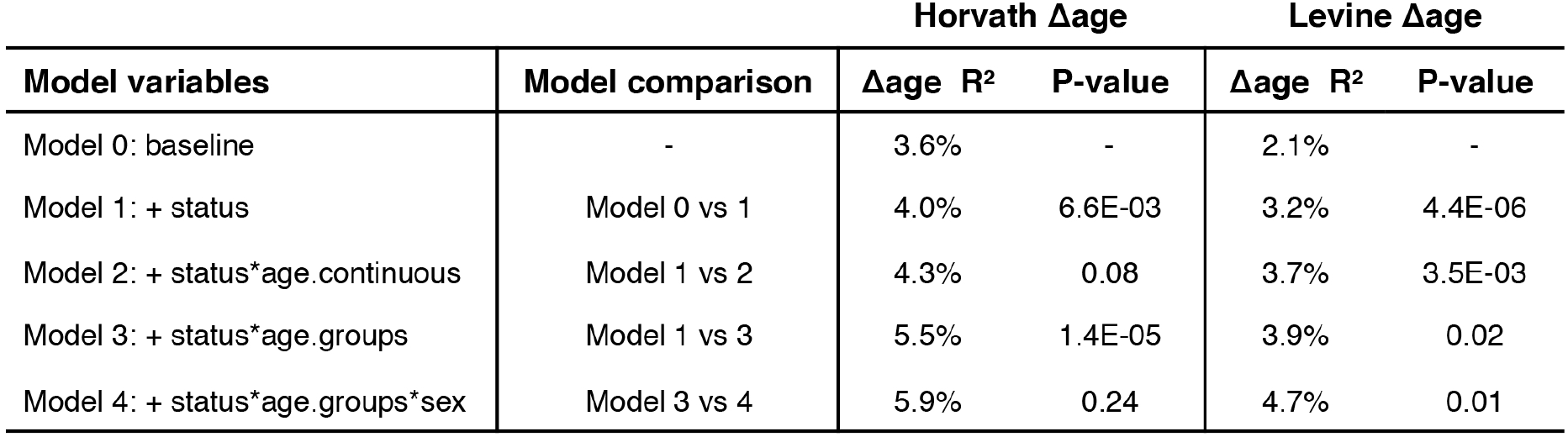
Age- and sex-specific effects significantly contribute to DNAm aging in schizophrenia. Shown are the contributions of interaction effects between disease status and age and sex on Δage. The baseline model corresponds to Δage ~ dataset + ethnicity + platform + age.continuous + sex. For other models, the variable(s) in addition to the baseline variables are shown with the corresponding variance explained (R2) in Δage. Interaction terms with chronological age are modeled as a continuous variable (age.continuous) or a categorical variable (age.groups). The latter uses previously defined decades. Model comparison is performed to assess if the contribution of an interaction term is significant compared to a model without that term. The chi-square test is used to test two models with corresponding p-value presented. The results of these analysis are shown for both the Horvath and Levine clock. P-values are corrected for the number of tests performed (2 clocks x 4 comparisons = 8).

### Estimating and mapping windows of differential aging in schizophrenia

As our categorical age groups in the previous analyses were chosen somewhat arbitrarily, we conducted an exploratory analysis to refine age-dependent aging effects to identify specific age windows that are associated with differential aging. We implemented a sliding window approach across chronological age, both in the full sample and within each sex separately. Using 5-year bins and sliding steps of 1 year, we tested cases versus age-matched controls and constructed a more precise picture of differential aging across chronological age in SCZ. At this finer resolution, we mapped changes in Δage to specific ages with different patterns between men and women. For intrinsic cellular age, we observe a deceleration effect during early adulthood from 29 years and younger across all samples, with the shift in differential aging occurring earlier in women (<25) (Figure 3C). For both men and women, we observe age deceleration in mid-forties and for women we also find age acceleration between 50-56 years (Figure 3C-E).

For phenotypic age, we mapped the age acceleration effect to 27 years and older across the whole sample with differences between the sexes (Figure 3F-H). In women, we find age acceleration between 25-29 years and from 36 years and older (Figure 3G). In men, we find age acceleration between 27-39 and 49-59 years (Figure 3H). More details on each age window and corresponding effect sizes are shown in Table S10. Thus far, our results show that DNAm aging, measured through the Horvath and Levine clock, is significantly different in SCZ and characterized by age-specific effects with some distinctions between the sexes, particularly for Levine Δage.

### DNAm aging affects SCZ above and beyond smoking and blood cell types

To investigate the effect of smoking and blood cell type composition, we use DNAm-based smoking and cell type estimations (see Methods) as a proxy to evaluate their contribution to DNAm aging in SCZ. While DNAm clocks, by design, will encapsulate such effects, quantifying the contributions of each factor increases interpretability and helps understand the factors contributing to the differential aging findings. We observe that blood cell type proportions explain significantly more variance in DNAm aging than DNAm smoking scores (Supplementary Results S2.1). Inclusion of DNAm smoking score and blood cell proportions as covariates in our main models explains part but not all of the observed disease effects (Table S11 and Figure S8-9). Using a penalized regression framework (Table S12), we show that Levine Δage independently contributes to the variance in disease status in women older than 36 above and beyond smoking scores and blood cell type proportions (Supplementary Results S2.2 and Figure S10). A significant proportion of the Horvath Δage effect on disease status is reduced by adjusting for smoking (Table S11). However, smoking is not associated with Horvath Δage in controls (Pearson r=0.01, P=0.95) nor in cases (Pearson r=−0.08, P=0.28) (Figure S11). As smoking covaries with SCZ disease status, it is difficult to distinguish these signals.

### Age deceleration by multi-tissue Horvath clock is not present in brain

We investigated DNAm aging in frontal cortex postmortem brain samples of 221 SCZ cases and 278 controls. The multi-tissue Horvath clock accurately predicts DNAm age in the brain as well (r=0.94, P < 2.2e-16). We, however, find no difference in DNAm aging between cases and controls (ß=−0.29, P=0.46) and no evidence of age-dependent aging. More details are shown in the Supplementary Results (S2.3).

### Phenotypic age acceleration is associated with SCZ polygenic risk in women

To further decipher the factors underlying the signal of differential aging in SCZ, we examined the possible role of SCZ polygenic risk, age at onset, and illness duration (Figure S12). We first focus on the phenotypic age acceleration in female SCZ patients of age 36 years and older, as these individuals showed the most consistent and pronounced aging effect. We find stronger age acceleration in cases with both low and high SCZ genetic risk (Table 2). More specifically, patients in the highest PRS1 tertile are predicted to be 4.30 years older in phenotypic age compared to controls (P=1.3E-05), patients with median range PRS1 are 1.89 years older (P=4.5E-02), and patients in the lowest quartile are 2.89 years older (P=2.8E-03). By permutation of PRS1 bins, we find that the effect in the highest PRS1 tertile is unlikely to occur by chance (P=0.024). For the association between Levine Δage and PRS1 to be most pronounced in the low and high tertile, is even less likely to happen by chance (P=0.006). At maximum, this group of women carrying high SCZ genetic risk have on average 7.03 higher phenotypic Δage (95% CI: 3.87-10.18; P=1.7E-05) (Figure 4A). We do not observe such an association in women age < 36 years, men with age > 36 years, nor across the whole dataset (Figure 4B and S13). Finally, by permuting the ranks of PRS1 within female cases >36 years, we find a mean maximum phenotypic Δage case-control difference of 3.69 years (95% CI: 1.26-6.12) across 1000 permutations, further demonstrating the significance of the observed maximum of +7.03 years phenotypic Δage difference. For age at onset and illness duration, we did not find significant association with Δage across partitioned bins (after permutation, P > 0.05) (Table 2).

**Table 2.** Integration of Levine Δage with PRS, age of onset, and illness duration in women in later adulthood. Analyses were performed using women >36 years of age. Only cases with available information were included in the analyses. Each phenotype was analyzed as both a continuous variable and as a categorical variable using equal tertiles from low to high bins. Mean values in cases for each phenotype are presented along with the association with Δage (β) and corresponding 95% confidence intervals and p-values. PRS1 = polygenic risk score PC1 (see Supplementary Information) scaled to mean zero with standard deviation of 1, AOO = age of onset, DUR = illness duration. Asterisk* indicates that significance (P < 0.05) by permutation analyses.

| Women: >36 | Controls | Cases | Mean value<br>in cases | $\beta$<br>(Levine Δage) | 95% CI | P |
| --- | --- | --- | --- | --- | --- | --- |
| **Polygenic risk** |  |  |  |  |  |  |
| All - no stratification | 227 | 149 | 0.35 | 3.02 | 1.76 — 4.27 | 3.1E-06 |
| PRS1 - continuous | - | 149 | 0.35 | 0.42 | -0.37 — 1.21 | 3E-01 |
| PRS1 - low | 227 | 50 | -0.68 | 2.89 | 1.00 — 4.77 | 2.8E-03 |
| PRS1 - mid | 227 | 50 | 0.34 | 1.89 | 0.05 — 3.73 | 4.5E-02 |
| PRS1 - high | 227 | 49 | 1.40 | 4.30 | 2.40 — 6.20 | **1.3E-05\*** |
| **Age of onset** |  |  |  |  |  |  |
| All - no stratification | 227 | 111 | 26.50 | 3.73 | 2.34 — 5.12 | 2.3E-07 |
| AOO - continuous | - | 111 | 26.5 | -0.08 | -0.21 — 0.05 | 2.2E-01 |
| AOO - early | 227 | 37 | 17.43 | 3.26 | 1.11 — 5.41 | 3.1E-03 |
| AOO - mid | 227 | 37 | 25.43 | 3.70 | 1.58 — 5.81 | 6.7E-04 |
| AOO - late | 227 | 37 | 36.62 | 4.24 | 2.09 — 6.40 | 1.3E-04 |
| **Illness duration** |  |  |  |  |  |  |
| All - no stratification | 227 | 111 | 23.37 | 3.73 | 2.34 — 5.12 | 2.3E-07 |
| DUR - continuous | - | 111 | 23.37 | 0.03 | -0.07 — 0.13 | 6.1E-01 |
| DUR - short | 227 | 37 | 10.76 | 3.90 | 1.76 — 6.03 | 3.9E-04 |
| DUR - mid | 227 | 37 | 23.33 | 2.91 | 0.78 — 5.05 | 7.7E-03 |
| DUR - long | 227 | 37 | 36.01 | 4.39 | 2.25 — 6.53 | 7.3E-05 |

**Figure 4.**
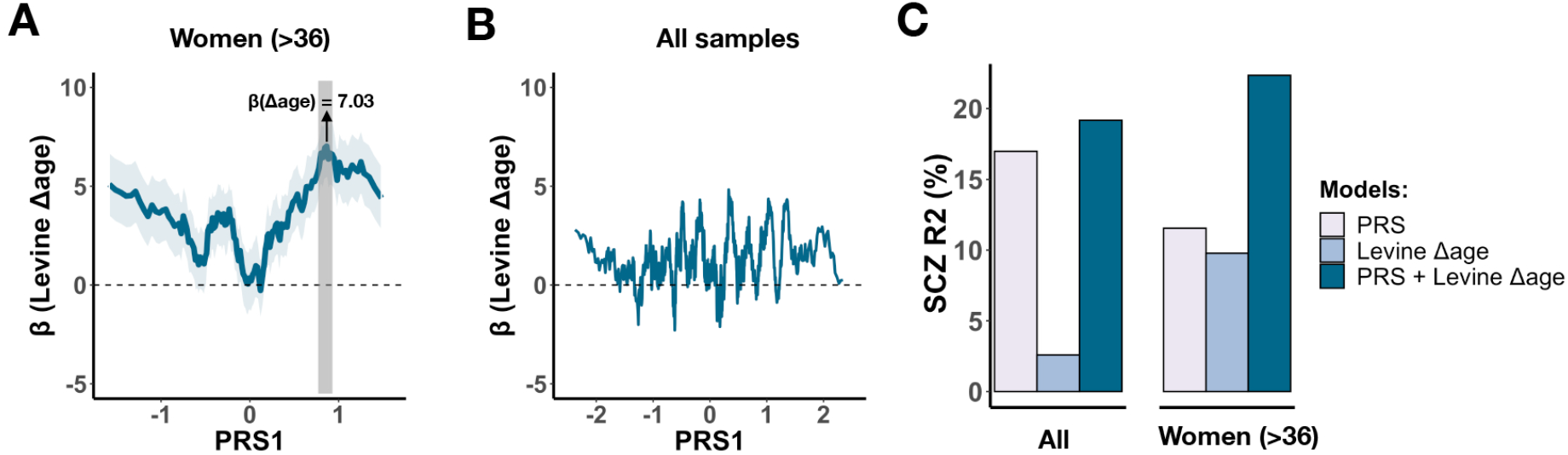
DNAm aging associates with SCZ PRS and additively contributes to SCZ disease status. (A) Using a sliding-window approach, Levine Δage difference between cases and controls are shown across bins of ranked PRS1. Each bin contains 20 cases and slides from low to high PRS1 per shifts of one sample. The estimated Δage difference compared to all female controls >36 years is shown for each sliding bin in blue with the standard error in shaded blue. The most significant bin is highlighted by the grey vertical bar. (B) A similar analysis but then across all samples. (C) The variance explained in schizophrenia disease status (y-axis) by SCZ PRS and Levine Δage shown for all samples (left) and for women in later adulthood (right). The estimates shown are derived on top of the effect of sex, ethnicity, batch, platform, and chronological age.

This is further confirmed when we integrated these two variables across PRS1 tertiles, demonstrating that the most pronounced differences in Δage are observed across PRS1 bins and not across the distribution of age at onset and illness duration in this subset of women (Figure S14).

We conducted a similar investigation on the observed intrinsic cellular age deceleration in all SCZ patients aged 29 years and younger but found no significant associations between Horvath Δage and PRS1, age at onset, or illness duration (Table S13 and Figure S15). While we did observe the strongest Horvath age deceleration in the high PRS1 tertile (β=−1.58, P=3.0E-03), this was not significant after permutation analysis (P>0.05). We did not analyse other identified age windows of differential aging as these either had too few individuals with genetic or phenotypic information available or more modest disease effects limiting any further stratification.

Finally, we assessed how Levine Δage and SCZ PRS1 compare in predicting SCZ disease status in our sample. Across the whole sample, PRS1 and Levine Δage explain 17.0% and 2.6% of the variance in disease status, respectively. Together, they explain 19.2%. In women in later adulthood, SCZ PRS1 and Levine Δage explain 11.5% and 9,8% independently and 22.4% jointly (Figure 4C).

## Discussion

We performed a large study of biological aging in schizophrenia using multiple epigenetic clocks based on whole blood DNA methylation data. We observe significant patterns of sex-specific and age-dependent DNAm aging in SCZ, a finding consistent across four European cohorts. The most significant differential aging pattern that we observe is in females ages 36 years and older in which we detect advanced *phenotypic age acceleration*, as measured by the Levine clock. We also observe *intrinsic cellular age deceleration* in SCZ patients during early adulthood, as measured by the Horvath clock. Phenotypic age acceleration in female patients is associated with a higher burden of SCZ polygenic risk. This high SCZ risk group displays accelerated aging of an average of +4.30 years compared to age-matched female controls. Phenotypic age and SCZ PRS furthermore contribute additively to SCZ and explain up to 22.4% of the variance in disease status. Our findings suggest that specific and identifiable patient groups are at increased mortality risk as measured by the Levine clock and warrant further research on DNAm clocks to examine its clinical relevance.

The Levine estimator was constructed by predicting a surrogate measure of phenotypic age, which is a weighted average of 10 clinical markers, including chronological age, albumin, creatinine, glucose and C-reactive protein levels, alkaline phosphatase and various blood cell related measures [17]. By design, the Levine estimator is a composite biomarker that strongly predicts mortality, in particular that of age-related diseases, such as cardiovascular-related phenotypes. A 1-year increase in phenotypic age is associated with a 9% increased risk of all-cause mortality and a 10% and 20% increase of cardiovascular disease and diabetes mortality risk, respectively [17,36]. Our findings of multiple year increase in phenotypic age in SCZ could thus imply an increased mortality in patients that is linked to cardiovascular disease, a previously well-established epidemiological observation [4,5,37]. A recent study however found that DNAm age acceleration only predicts mortality in SCZ cases without pre-existing cancer using the Hannum clock [38]. They did not find such evidence using the Levine clock. The smaller sample size and predominantly male cohort may have reduced the predictive power of the study. Our findings warrant a more focused and larger study of DNAm aging in female patients in later adulthood, preferably stratified by SCZ genetic risk. Our results align well with the observation that patients with SCZ, particularly women, are reported to be at high mortality risk due to cardiovascular disease and diabetes [5,39,40]. Assuming that cardiovascular risk is modifiable in SCZ [41], phenotypic age could serve as a potential biomarker to identify at-risk individuals and in this way help with disease management and improvement of life-expectancy.

In contrast to *age acceleration* in phenotypic age, we observe *age deceleration* in intrinsic cellular age (i.e. the Horvath DNAm age), an effect that is most pronounced in patients age 29 and younger. Unlike the association findings in females, we did not observe clear patterns with genetic and phenotypic variables that could help to further decipher the signal. Horvath Δage furthermore showed strong age-specific effects but less clear sex-specific effects. We did not observe *age deceleration* in postmortem brain samples of the human cortex, indicating that the observed aging signal in SCZ may be blood-specific. Horvath DNAm aging has been shown to be associated with molecular processes of development and cell differentiation [15,35], including through blood-based DNAm age measures in human (neuro)developmental phenotypes [42,43]. Our findings may indicate that patients diagnosed with SCZ in this age group show evidence of delayed or deficient development and that this is detectable in blood through the multi-tissue Horvath clock. This however remains speculative and future work is needed to further dissect how blood-based *Horvath age deceleration* is associated with SCZ.

While we did observe aging effects with the Horvath and Levine clock, we did not with the Hannum clock. The Hannum clock is less predictive of age acceleration effects on mortality risk than the Levine clock [17], which could explain the lack of findings in our analyses. The Hannum estimator furthermore cannot be used on first generation 27K DNA methylation arrays which reduced the sample size of this study with 30% and may have impacted the statistical power of these specific analyses. This highlights the benefits of designing methods that are inclusive to all platforms, so all data, both old and new, can be leveraged.

After publication of the preprint of our manuscript [44], Higging-Cheng et al. also reported significant DNAm alterations in SCZ [45]. This smaller study included 567 SCZ cases and 594 non-psychiatric controls with most of the sample (UK and SCT cohorts) also included in our study. Similar to our finding of 1.53 years of phenotypic age acceleration in schizophrenia cases, they report a 1.4- to 1.9-year increase in Δage in SCZ cases compared to controls. In addition, using GrimAge, a newly trained DNAm mortality clock [46], they observe age acceleration of 2.5- to 5.8-years. Unlike phenotypic age acceleration, this increase is largely driven by smoking effects. Similar to our work, this work highlights the value of analysing multiple clocks in conjunction and again suggesting that distinct biological processes of aging are altered in SCZ. In addition to the larger sample size, there are other key differences between our study and Higgins-Cheng et al. First, we performed detailed phenotypic analyses including explicit modelling of age and sex-specific effects. Second, methodically, we performed meta-analyses across cohorts as opposed to individual analyses per cohort. This approach, combined with multiple testing correction, is robust to cohort-specific artefacts in the data. Third, we integrated DNAm age with SCZ polygenic risk. Our PRS analyses yielded important insights into specific patient groups that could be at higher risk of all-cause mortality and that DNAm Δage and SCZ polygenic risk contribute additively to the illness. The latter suggests that combining genetic and epigenetic predictors can augment downstream prediction of outcomes in SCZ, similarly to what was recently shown for BMI [47].

A systematic review of aging biomarkers found that less than a quarter of studies explored an interaction effect or statistically compared the regression slope between groups in SCZ [14]. Our findings of sex-specific and age-dependent DNAm aging support their recommendations to specifically examine interaction effects with age and sex in aging studies but also more general in epigenetic studies of SCZ, such as epigenome-wide association studies. Future work should also be extended to integrate nonlinear models to fully capture the complex relationship between DNAm aging and clinically relevant variables across the lifespan of patients. These models will help validate and further refine the most relevant age intervals.

A limitation of the study is the cross-sectional design of the cohorts used. While we do find an association with SCZ polygenic risk, dissecting cause-and-effect relationships remains challenging. Independent replication studies are needed, preferably using longitudinal prospective cohorts with genomic data and information on symptom recurrence and severity, comorbidities and other phenotype-related variables. These studies can assess the clinical relevance of DNAm aging in SCZ above and beyond other known health risk factors and disease biomarkers, such as medication use. An urgent open question remains whether DNAm age signatures are modifiable with regards to clinical and lifestyle factors associated with SCZ. Improvement of existing methodology and/or development of new DNAm age biomarkers [48,49] may in addition help to better study differential aging in SCZ and related disorders with increased mortality. Combining blood-based DNAm age with that of other aging profiles, such as MRI-based brain age [50], may further advance our understanding of aging and SCZ disease progression, including the increased mortality [51]. Finally, our findings support an integrative strategy with polygenic disease risk to improve clinical utilization.

Schizophrenia, like other mental illnesses, are associated with a wide-range of subsequent chronic physical conditions, including many age-related diseases [52]. While health and life expectancy of the general population continues to improve, the mortality disparity between patients with schizophrenia and those unaffected continues to increase [9,10,53,54]. As the burden of age-related diseases continues to rise, early detection and subsequent opportunities for interventions before disabilities and co-morbidities become established will be important [1,2]. Molecular biomarkers of aging, such as DNAm clocks, are now emerging as candidate tools for screening and intervention. Taken together, this study strengthens the need for more research on DNA methylation aging in SCZ, a population vulnerable to age-related diseases and excess mortality.

## Material and Methods

### Cohort and sample description

Details of samples included in this study can be found in the Supplementary Information. Briefly, unrelated patients with SCZ and ancestry-matched non-psychiatric controls from four cohorts of European ancestry were included; the Netherlands (N=1,116), Scotland (N=847), Sweden (N=96), and the United Kingdom (N=675). Cases were selected on the basis of a clinical diagnosis of SCZ using the Diagnostic and Statistical Manual for Mental Disorders (DSM-IV), Research Diagnostic Criteria (RDC), or the International Classification of Diseases 10 (ICD10). Controls were unaffected subjects without a history of any major psychiatric disorder. Whole blood DNAm data was available for a total of 2,707 samples (1,399 cases and 1,308 controls; Table S1).

### Genome-wide DNA methylation profiling and data processing

To quantify DNA methylation, DNA was extracted from whole blood and bisulfite converted for hybridization to the Illumina Infinium Human Methylation Beadchip. Samples were assayed with either the 27K or 450K beadchip, which contain 27,578 and 485,512 probes that interrogate CpG sites across the genome, respectively. For each platform, data processing pipelines were implemented, which includes background correction, color channel and probe type correction, and normalization of the data, to minimize the effect of technical variation on the final beta values. Samples with more than 5% of probes detected at P > 0.05 were excluded from further analyses (n=13). Full details are described in the supplementary methods.

### DNAm-based estimation of biological age

To compute blood-based DNAm age estimates, processed beta values were used as input to the Hannum[16], Horvath[15], and Levine [17] DNAm clock. These DNAm age estimators use a set of CpGs that are selected via an optimization algorithm to collectively minimize the error associated with estimating chronological age (Supplementary Information). Horvath DNAm age estimates were calculated using R scripts from the Horvath DNA Methylation Calculator (https://dnamage.genetics.ucla.edu). Hannum and Levine estimates were obtained by using the reported set of probes with corresponding regression weights. We define Δage by subtracting chronological age at the time of the blood draw from the predicted DNAm age.

### Statistical analyses

To investigate epigenetic aging differences in SCZ, we first removed samples with discrepant phenotypic sex and predicted sex based on DNAm data (n=9), as well as samples with missing chronological age data (n=237), bipolar disorder diagnosis (n=26), and duplicate samples (n=126). For each epigenetic clock, we regressed Δage on technical principal components (PCs), using the first components that cumulatively explain >90% of variation in intensity values of control probes, and added the residuals to mean(Δage) to generate a measure in the same units as Δage that is adjusted for technical variation (Δage-adjusted). We used the adjusted value for subsequent analyses and refer to it as Δage.

As association analyses of DNAm age between groups are sensitive to the distribution of chronological age, particularly at older ages, any case older than the oldest control was excluded from each cohort (n = 5 for NLD, 16 for SCT, 4 for SWD, and 1 for UK). Chronological age was furthermore included as a covariate in all analyses, as recommended[55]. To minimize the effect of outlying samples, we excluded samples >3SD from mean Δage across cohorts (ranging from n=13 to 16 for the three clocks). These are samples for which DNAm age diverged substantially from chronological age, which are likely artifacts.

For each clock and each cohort, we implemented a multivariable regression model predicting Δage as a function of schizophrenia status, sex, and age. For the Dutch cohort, batch and array platform were also included as covariates, as this cohort consists of multiple datasets from both the 27K and 450K platform. For each clock, regression coefficients with corresponding standard errors for each of the four cohorts were then supplied to the rma() function of the metafor package[56] in R to fit a meta-analytic fixed-effect model with inverse-variance weights and obtain an overall effect size and test statistic. To quantify the significance of age- and sex-specific effects, we determined the contribution of interaction effects on top of the main disease effect. We first combined all cohorts to maintain necessary sample sizes across age and sex groups. Age groups were defined by grouping samples by decades with ages 18 and 19 included in the first decade (18-30, 31-40, etc.). To quantify the gain in variance explained in Δage, models with the interaction term were compared to a baseline model without the interaction term. For each analysis, statistical significance was determined using Bonferroni correction, i.e. P < 0.05 / number of tests.

### SCZ polygenic risk quantification

Polygenic risk scores (PRS) were obtained from analyses of the SCZ GWAS conducted by Psychiatric Genomics Consortium (PGC)[57]. Using a leave one out approach, weights were generated in a training dataset based on all samples minus the target cohort in which the PRS were calculated. For each individual, weighted single nucleotide polymorphisms (SNPs) were summed to a genetic risk score that represents a quantitative and normally distributed measure of SNP-based SCZ genetic risk. To reduce between cohort-variation and maximize statistical power, we used a previously developed analytical strategy that uses principal component analysis (PCA) to concentrate disease risk across PRSs of ten GWAS p-value thresholds into the first principal component (PRS1)[58] (Supplementary Information). PRS1 explains 70.7% of the variance in risk scores and 19.9% of the variance in SCZ status, which is more than any of the original p-value thresholds (4.9-17.4%). The other PCs had no explanatory value in disease status (mean R^2^ = 0.0%), which means that PRS1 captures the majority of SNP-based SCZ polygenic risk. PRS1 was generated for 1,933 individuals, 853 cases and 1080 controls, and modelled as both a quantitative and categorical variable to predict Δage.

### Defining age at onset and illness duration

Age at onset is defined as the earliest reported age of psychotic symptoms or by the Operational Criteria Checklist (OPCRIT), depending on the cohort. This data is available for a subset of cases (N = 710) across the Dutch, Scottish, and UK cohorts. Illness duration is defined as the time between age at onset and blood collection. A more detailed description of each cohort’s definition is available in the Supplementary Information.

### DNA methylation-based smoking scores and blood cell type proportions

Smoking scores and blood cell type proportions were estimated from the data (see Supplementary Methods) and used as a proxy to further decompose differential aging effects.

### Estimating the contribution of differential aging in schizophrenia

Using a multivariable logistic regression model for disease status, we fitted batch, cohort, DNAm smoking score, DNAm blood cell type proportions, and Δage as explanatory variables. We first performed a variable reduction step to select the most contributing variables to disease status by use of a regularized logistic regression using the glmnet() function in R (“glmnet” package, v2.13)[59]. Alpha was set to “1” (Lasso) and the lambda parameter estimated at the optimal value that minimizes the cross-validation prediction error rate using the cv.glmnet() function. For each selected variable, we then report the variance explained in SCZ status (glm, family = “binomial”) for both the individual variable as well as adjusted for all other selected variables using the NagelkerkeR2() function in the “fmsb” package (v 0.6.3). The significance of each variable to their contribution was computed by comparing the model with and without the variable of interest using the likelihood ratio test of the anova() function.

## Declarations

### Ethics approval and consent to participate

All cases and controls included in this study gave informed consent. Dutch (NLD) cohorts - ethical approval was provided by local ethics committees; University College London (UK) cohort - ethical approval was provided by National Health Service multicentre and local research ethics; Aberdeen (SCT) cohort - was provided by both local and multiregional academic ethical committees. Sweden (SWD) cohort - ethical permission was provided by the Karolinska Institutet Ethical Review Committee in Stockholm, Sweden.

### Availability of data and materials

The datasets used are available on the NCBI Gene Expression Omnibus (GEO) data repository or through the principal investigator of each cohort. See Table S2 and S3 for an overview and corresponding accession series numbers. See Table S4 for sample information, including individual DNAm age estimates.

### Authors’ contributions

APSO and RAO conceived of the study. APSO performed data analyses and primary writing of the manuscript. LMOL, SH, and RAO advised on the work and co-wrote. RAO oversaw the work. RAO, RSK, EH, ED, DSC, NJB, AM, JM, JG and PFS provided access to data of cohorts included in the study. All authors read, gave input on, and approved the final manuscript.

## Acknowledgements

We thank Dr. Hannah Elliott (University of Bristol MRC Integrative Epidemiology Unit) for providing code to calculate DNA methylation smoking scores. We thank all study participants for their participation in each of the respective cohorts.

