## Supplementary Figures for "Epigenetic age is accelerated in schizophrenia with age- and sex-specific effects and associated with polygenic disease risk"

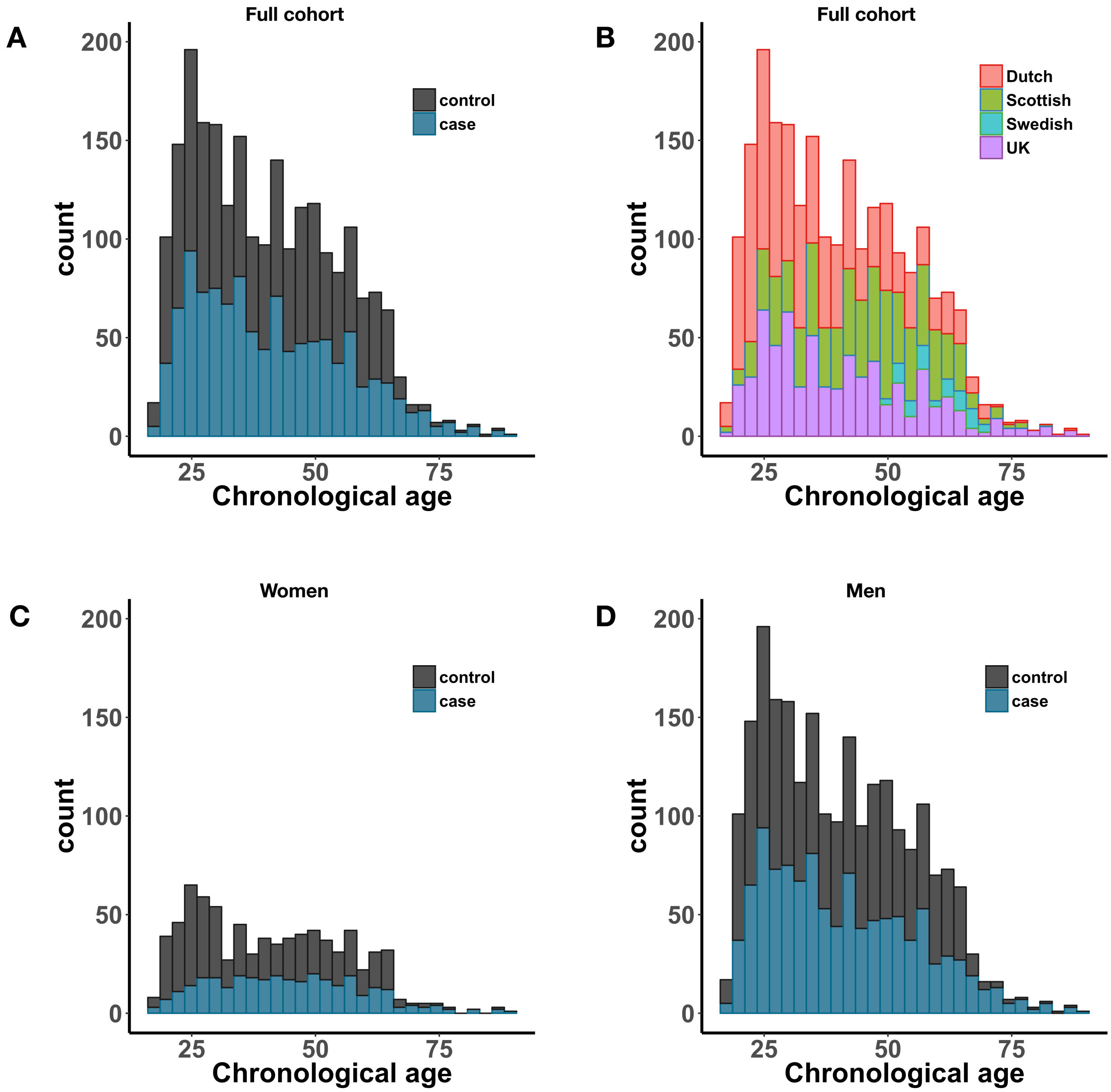


**Figure S1. Sample distribution across chronological age.** (A) Full sample colored by disease status. (B) Full sample colored by ethnicity. Women (C) and men (D) colored by disease status.

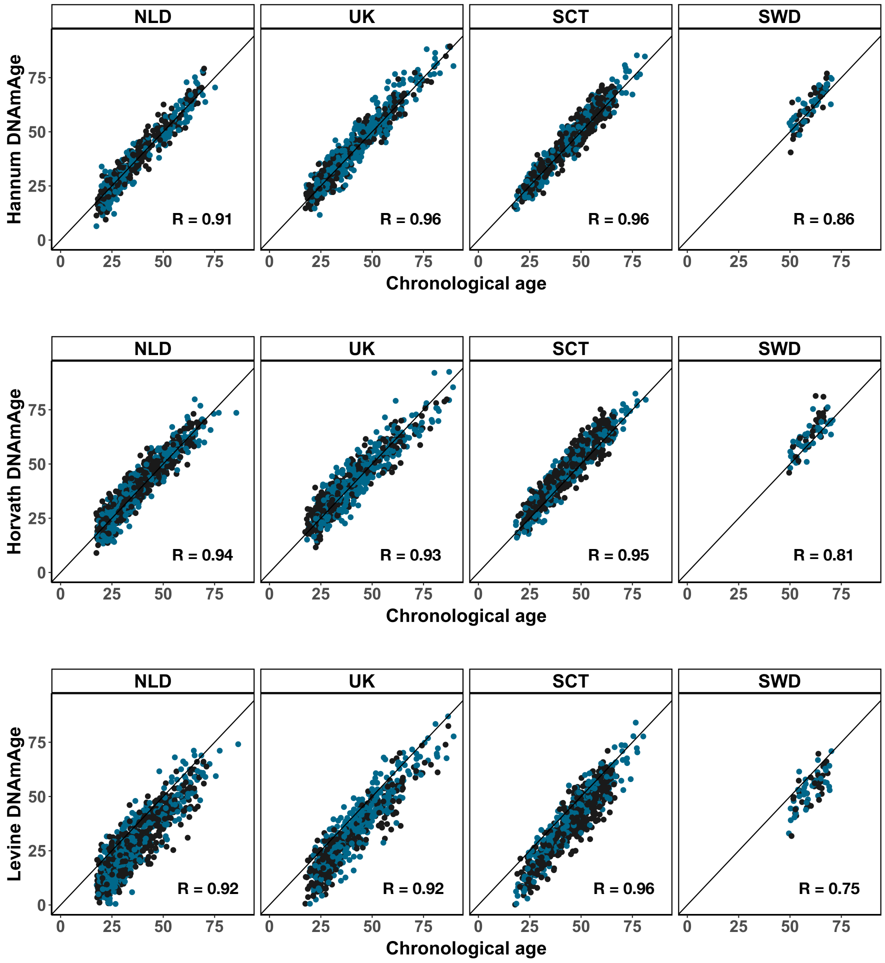


**Figure S2. Correlations between DNAm age and chronological age by ethnicity.** Shown are correlations for the Hannum (top), Horvath (middle), and Levine clock (bottom) across cohorts.

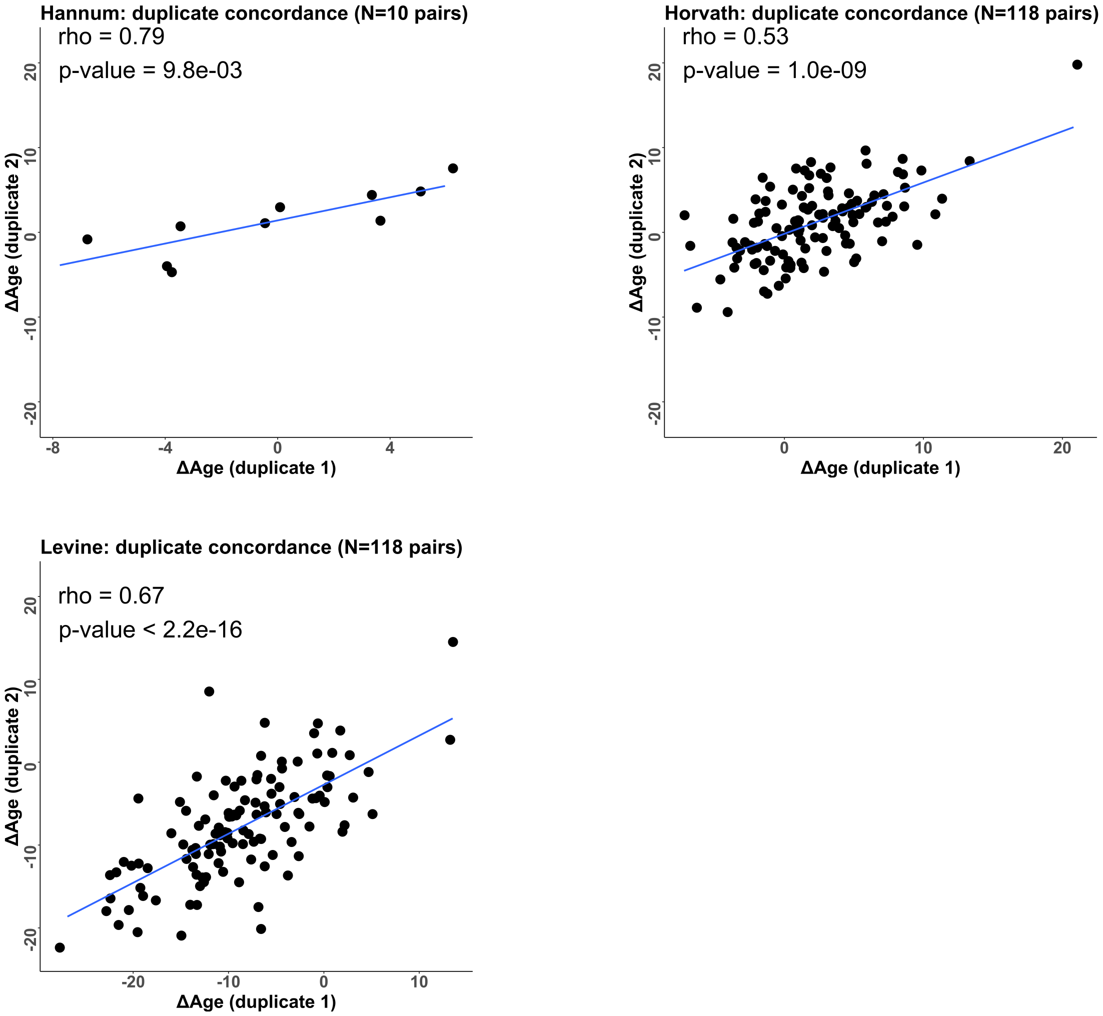


**Figure S3. Sample duplicate pair concordance for DNAm age estimates of the Horvath and Levine clock.** Using Δage across 118 duplicate pairs, the concordance between pairs is shown for the Horvath (top-left) and Levine (top-right) plot. For the Hannum clock (bottom), only duplicates with both samples on the 450K array (N=10) could be used.

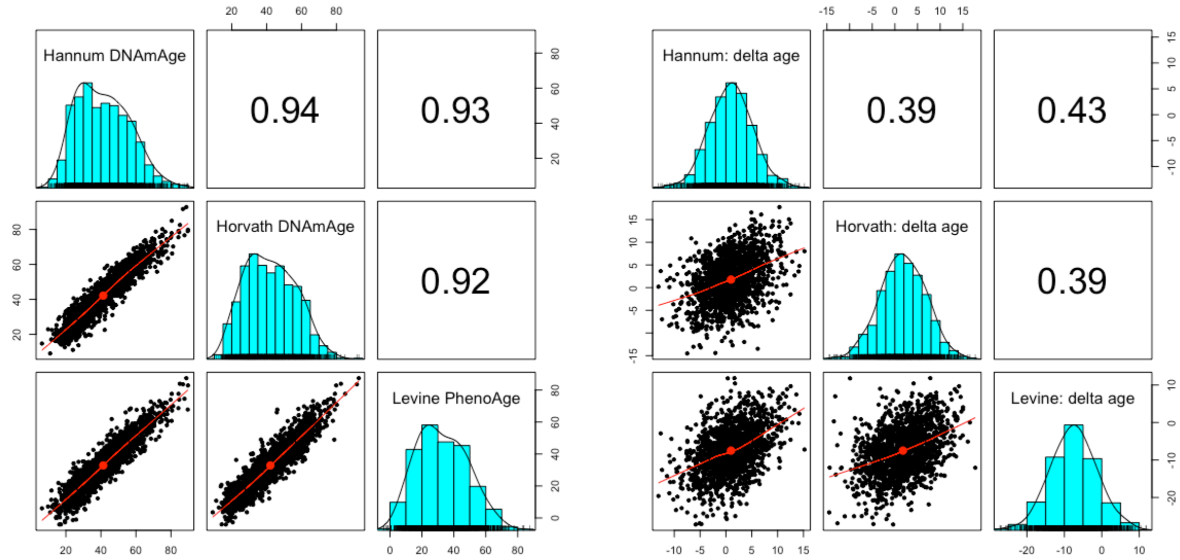


**Figure S4. Correlation structure across clocks highlights that they capture both shared and distinct aspects of aging.** Shown are pair-wise scatter plots below the diagonal, histograms on the diagonal, and the Pearson correlation above the diagonal for DNAm age (left) and Δage (right) across the three clocks.

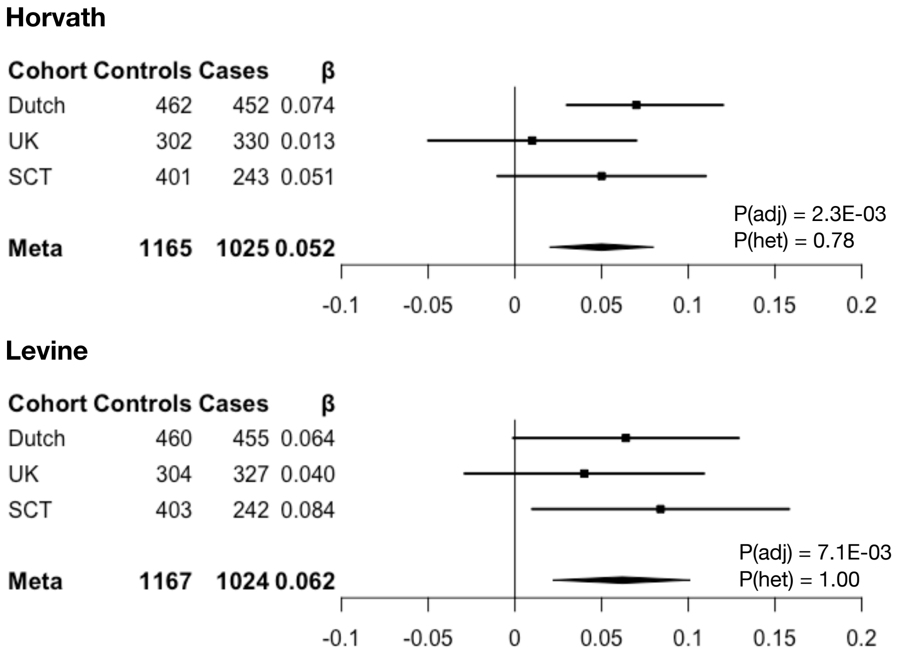


**Figure S5. Differential DNAm aging across cohorts.** Shown are results for modeling the interaction between disease status and chronological to estimate Δage differences between cases and controls conditional on age. For each estimator - Hannum (top), Horvath (middle), Levine (bottom) - number of cases and controls, and meta-analytic effect size (β) and adjusted p-value (Padj) are presented. See Table S4 for more details on results and corresponding statistics. The Swedish cohort was excluded from this analysis as it has only a limited spread in age (i.e. 50-70 years).

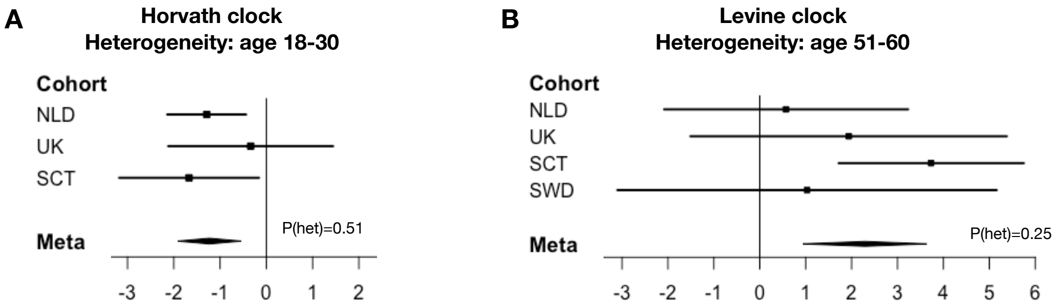


**Figure S6. Heterogeneity of aging effects within age groups across cohorts.** Shown are results for Δage differences between cases and controls in specific age groups for the Horvath (left) and Levine (right) clock. Each forest plots show a significant meta-analytic effect size and the p-value of Cochran's heterogeneity test (Phet). See Table S7 and S8 for more details on results and corresponding statistics. The Swedish cohort was excluded from the left plot as it has only a limited spread in age (i.e. 50-70 years).

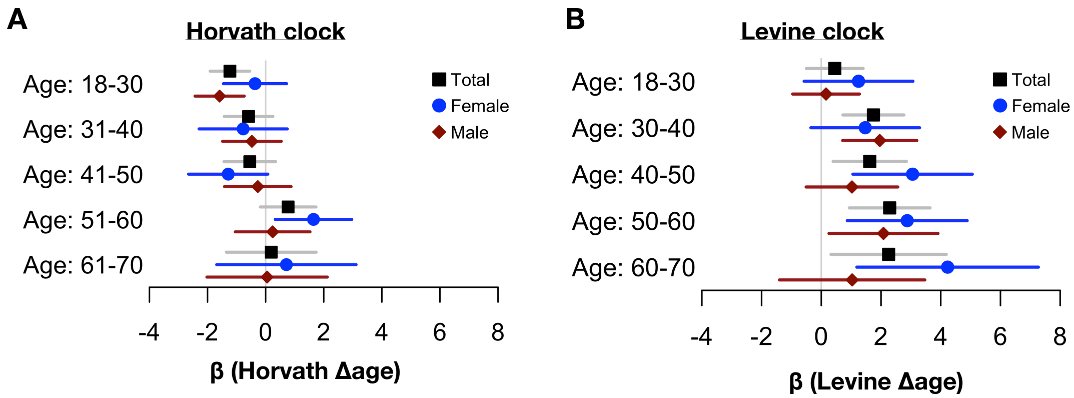


**Figure S7. Sex-stratified differential aging by age groups in schizophrenia.** Shown are Δage differences between cases and controls across age groups stratified by sex for the Horvath (A) and Levine clock (B). Results for women and men are presented in blue and red, respectively. The effects in the total sample are displayed in black.

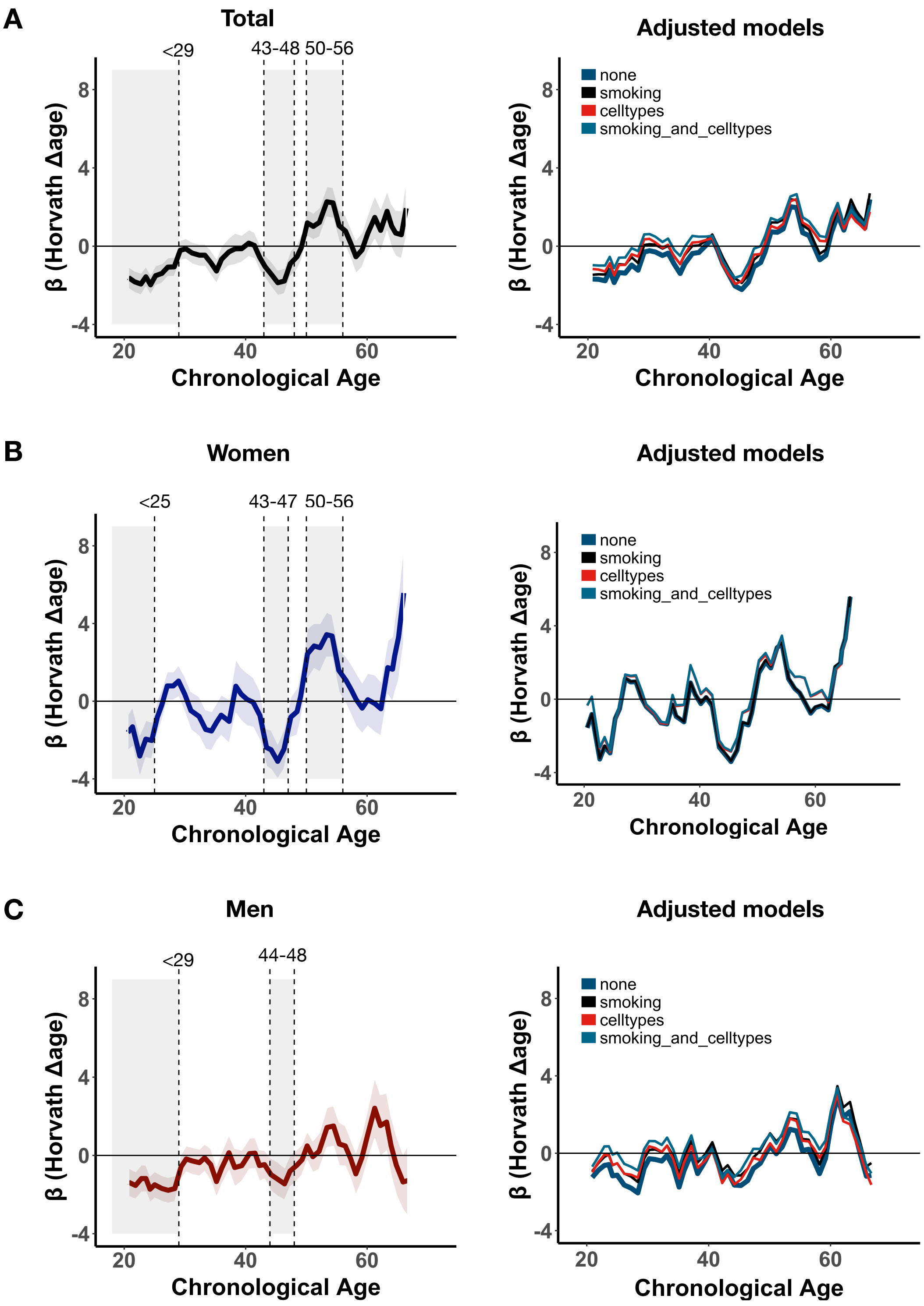


**Figure S8. Horvath differential aging across chronological age adjusted for DNAm smoking and cell type proportions .** Sliding age-windows, using 5-year bins with steps of 1-year, were used to estimate differential aging (β) at finer resolution across the range of chronological age. Graphs on the left show results without adjustment for smoking and cell types. Graphs on the right show results with adjustment for smoking and cell types. A) total sample size, B) women only, C) men only. For the right graphs only 450K samples were included, as DNAm smoking and cell types estimates cannot be calculated for 27K samples. These analyses therefore include less samples.

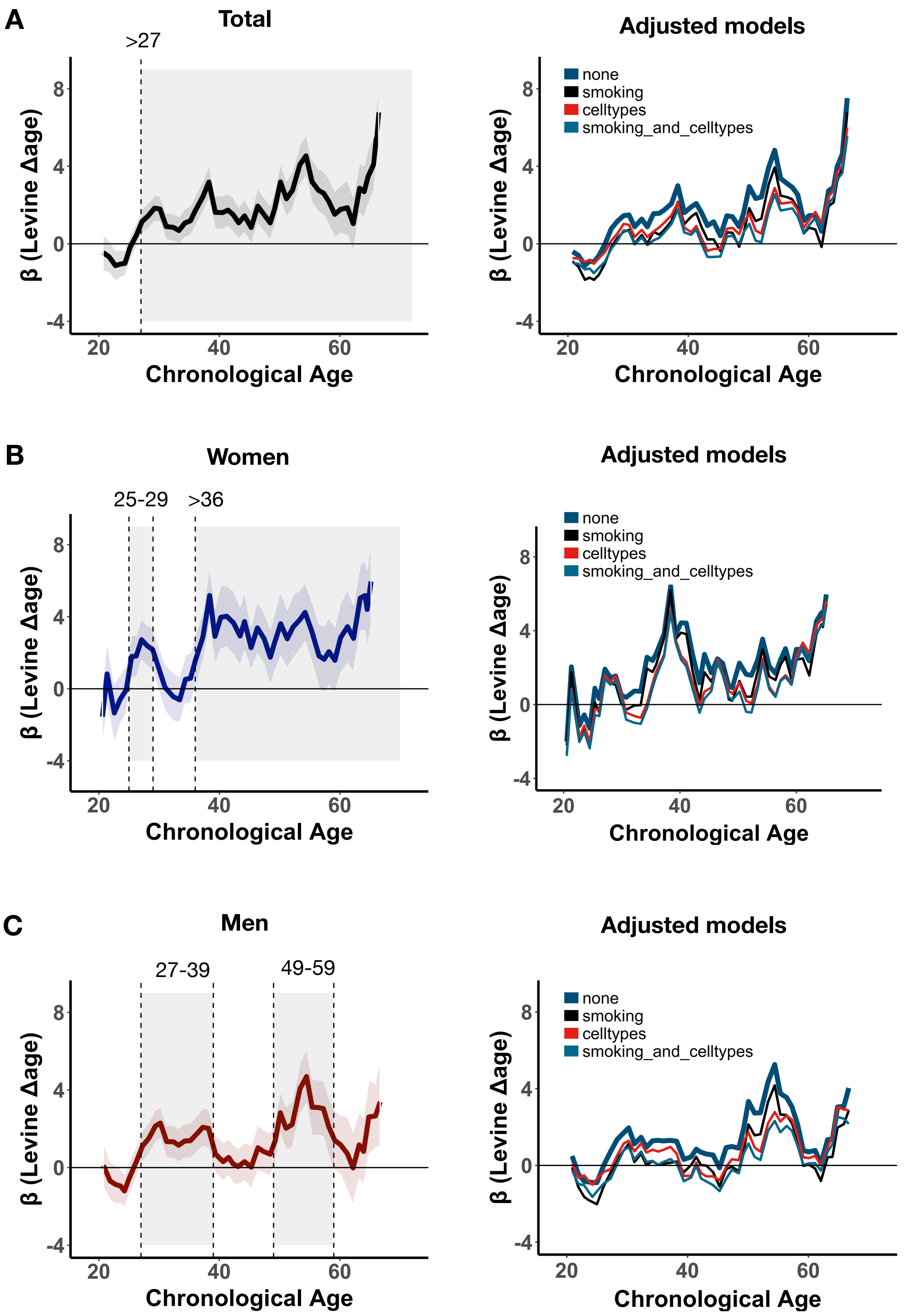


**Figure S9. Levine differential aging across chronological age adjusted for DNAm smoking and cell type proportions .** Sliding age-windows, using 5-year bins with steps of 1-year, were used to estimate differential aging (β) at finer resolution across the range of chronological age. Graphs on the left show results without adjustment for smoking and cell types. Graphs on the right show results with adjustment for smoking and cell types. A) total sample size, B) women only, C) men only. For the right graphs only 450K samples were included, as DNAm smoking and cell types estimates cannot be calculated for 27K samples. These analyses therefore include less samples.

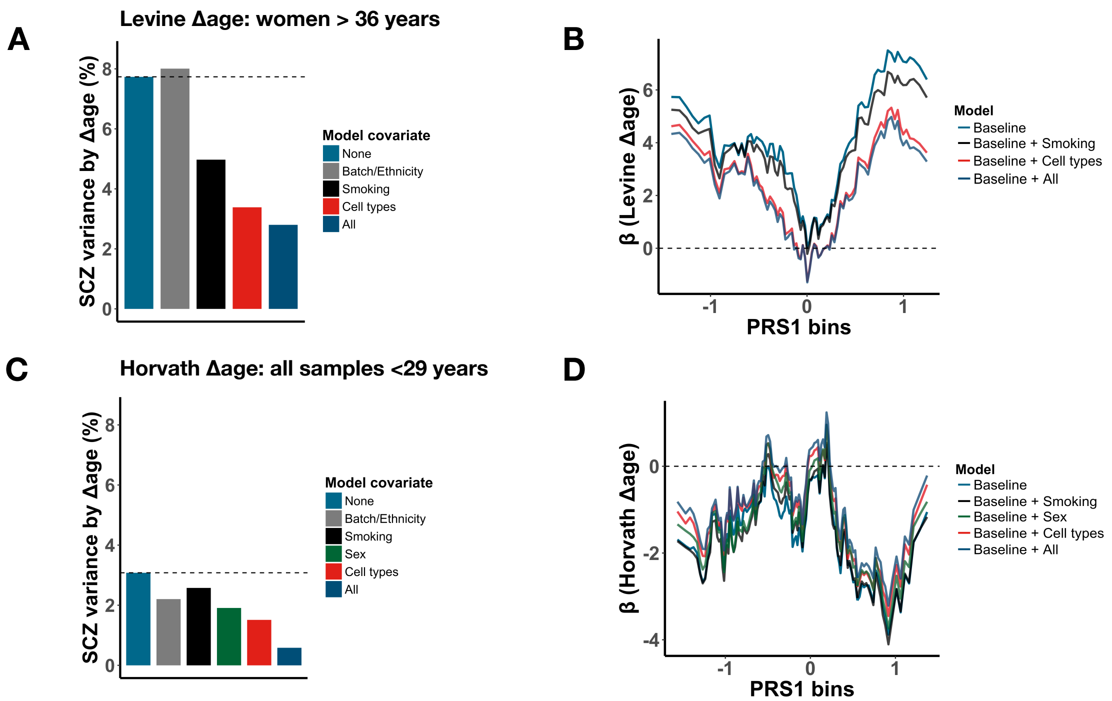


**Figure S10. Smoking and blood cell type composition contribute in part to DNAm aging.** Presented are the results of a sensitivity analysis of DNAm-based estimated smoking score and blood cell type proportions in the 450K subsample of the cohort. (A/C) The proportion of schizophrenia variance explained by Δage after adjustment of various variables that are significantly associated with disease status. The "All" model presents the variance explained by Δage independent from all other variables. (B/D) Δage effect size is shown across bins (N = 20 cases/bin) of ranked PRS1 (unit = SD) for several models that adjust for covariates. The baseline model represents the effect of Δage adjusted for batch, ethnicity and chronological age. Results are shown for Levine Δage women > 36 years, 99 cases and 181 controls (C) and Horvath Δage all samples < 29 years, 141 cases and 238 controls (D) separately.

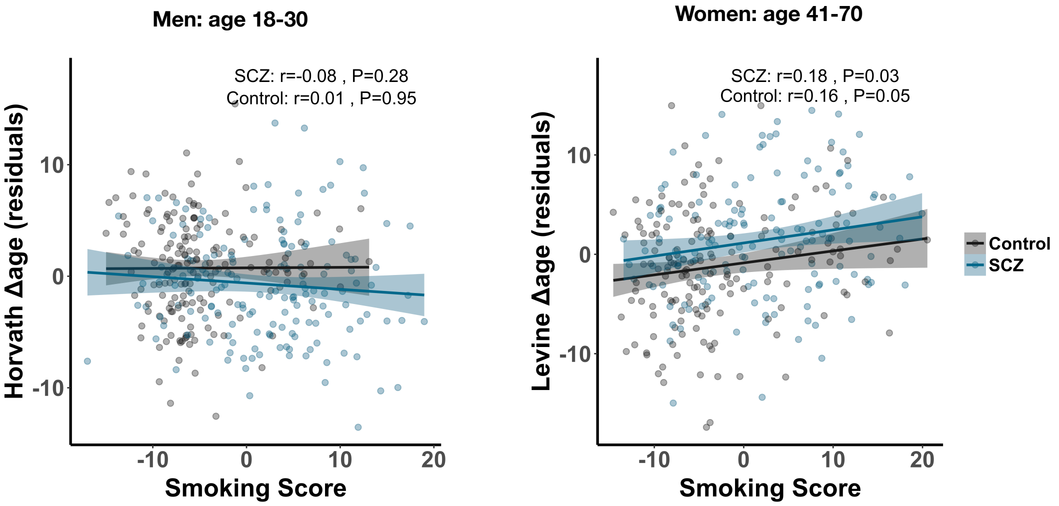


**Figure S11. Smoking has only minimal effects on DNAm aging.** Shown is the relationship between Δage and DNAm-based smoking score for men (left plot; age 18-30, case = 165, control = 163) and women (right plot; age 41-70, case=144, control=159). DNAm Δage was first regressed on batch, ethnicity and chronological age. The Pearson correlations and corresponding p-values are shown on top of each plot. Cases and controls are presented in blue and black, respectively.

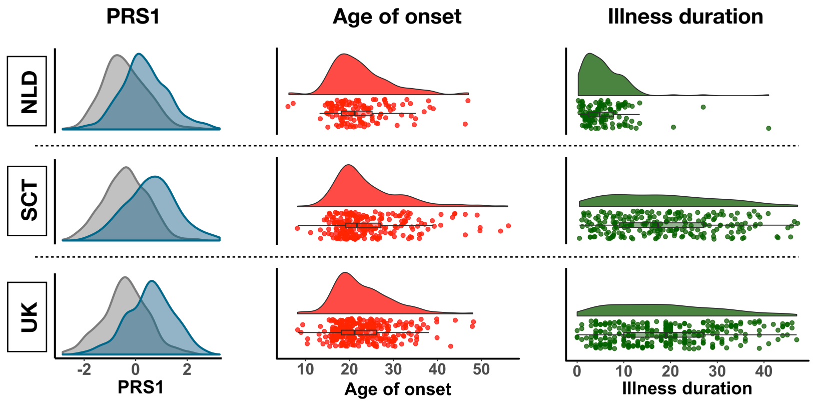


**Figure S12. Variable distribution of PRS, age of presentation, and illness duration.** The distribution of SCZ PRS1 score (left; cases in blue), age of presentation (middle), and illness duration (right) for each of the three cohorts with available information.

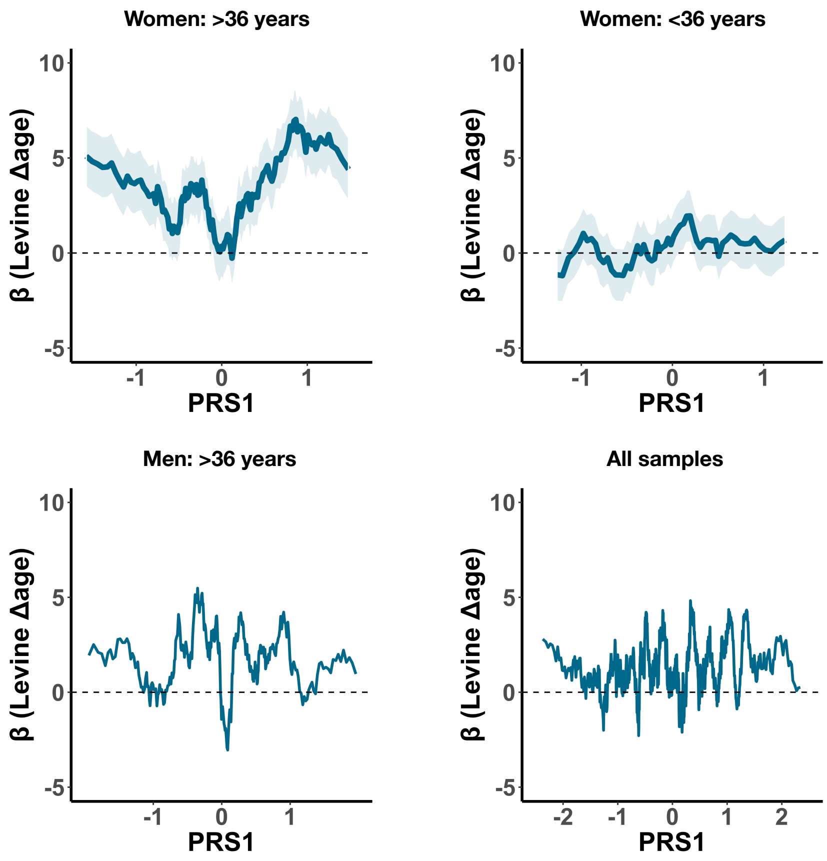


**Figure S13. The association between DNAm aging and PRS1 is unique to women > 36.** Using a sliding-window approach, Δage difference between cases and controls are shown across bins of ranked PRS1. Each bin contains 20 cases and slides from low to high PRS1 per shifts of one sample. The Levine Δage effect in each bin is shown in blue with the standard error in shaded blue. If the standard error is not shown, it was dropped to increase visual clarity. Results are presented for women > 36 (top left), women < 36 (top right), men > 36 (bottom left), and all samples (bottom right).

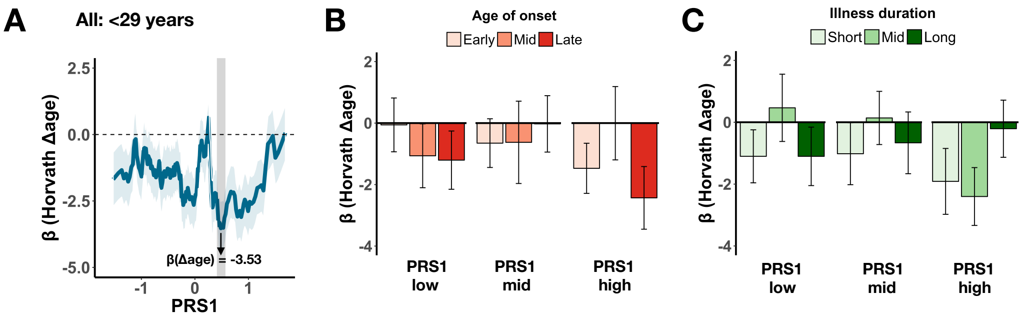


**Figure S14. Integration of DNAm aging with PRS, age of presentation, and illness duration across identified age intervals.** (A) Using a sliding-window approach, Horvath Δage difference between cases and controls are shown across bins of ranked PRS1. Each bin contains 20 cases and slides from low to high PRS1 per shifts of one sample. The estimated Δage difference compared to all male controls < 29 years is shown for each sliding bin in blue with the standard error in shaded blue. The most significant bin is highlighted by the grey vertical bar. (B ) DNAm aging effects stratified by PRS1 and age of onset. (C) DNAm aging effects stratified by PRS1 and illness duration.

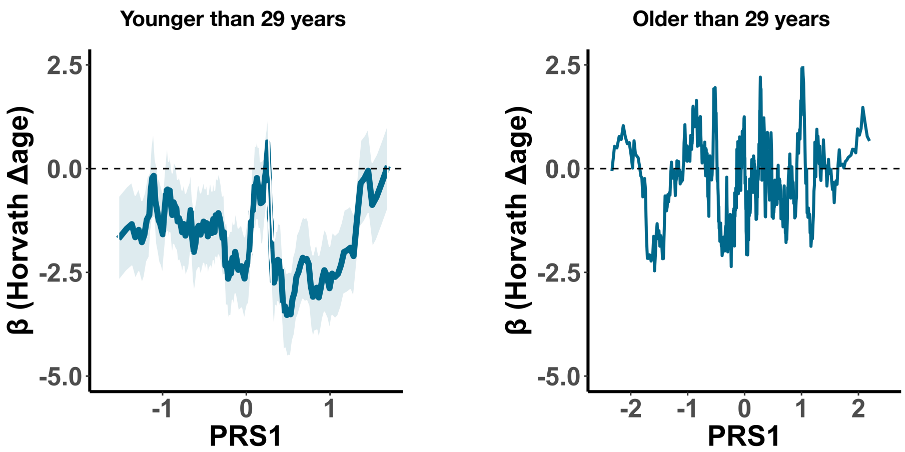


**Figure S15. The association between Horvath Δage and PRS1 is more pronounced <29 years.** Using a sliding-window approach, Δage difference between cases and controls are shown across bins of ranked PRS1. Each bin contains 20 cases and slides from low to high PRS1 per shifts of one sample. The Horvath Δage effect in each bin is shown in blue with the standard error in shaded blue. If the standard error is not shown, it was dropped to increase visual clarity. Results are presented for all samples < 29 (left) all samples > 29 years (right).
