## Supplemental Tables for "Epigenetic age is accelerated in schizophrenia with age- and sex-specific effects and associated with polygenic disease risk"

| | NLD | SCT | SWD | UK | Total |
|---|---|---|---|---|---|
| **Samples (N)** | 926 | 665 | 69 | 636 | 2,296 |
| **Cases** | 461 | 260 | 37 | 332 | 1,090 |
| **Controls** | 465 | 405 | 32 | 304 | 1,206 |
| **Age (years)** | 35.8 (13.2) | 44.6 (12.9) | 59.7 (5.94) | 40.4 (15.0) | 40.3 (14.4) |
| **Cases** | 35.5 (13.3) | 44.2 (14.1) | 59.6 (6.30) | 43.7 (14.6) | 40.9 (14.8) |
| **Controls** | 36.1 (13.2) | 44.9 (12.2) | 59.9 (5.58) | 36.8 (14.7) | 39.8 (14.1) |
| **Females** | 309 (33.4%) | 185 (27.8%) | 39 (56.5%) | 259 (40.7%) | 792 (34.5%) |
| **Cases** | 122 (26.5%) | 83 (31.9%) | 20 (54.1%) | 90 (27.1%) | 315 (28.9%) |
| **Controls** | 187 (40.2%) | 102 (25.2%) | 19 (59.4%) | 169 (55.6%) | 477 (39.6%) |
| **Hannum DNAm age** | 37.9 (15.4) | 46.0 (14.1) | 61.7 (8.37) | 41.6 (15.7) | 43.1 (15.6) |
| **Hannum Δage** | 0.0 (4.7) | 1.4 (4.2) | 2.0 (4.4) | 1.2 (4.7) | 1.0 (4.6) |
| **Horvath DNAm age** | 37.0 (14.0) | 47.6 (13.5) | 62.0 (7.7) | 41.2 (14.5) | 42.0 (15.0) |
| **Horvath Δage** | 1.2 (4.8) | 3.0 (4.7) | 3.2 (4.5) | 0.9 (5.7) | 1.7 (5.12) |
| **Levine DNAm age** | 27.5 (15.1) | 37.6 (15.4) | 52.9 (8.1) | 32.6 (16.5) | 32.6 (16.4) |
| **Levine Δage** | -8.33 (6.7) | -7.0 (6.2) | -6.8 (5.4) | -7.8 (6.6) | -7.7 (6.5) |

**Table S1. Sample characteristics and DNA methylation age estimates across cohorts.** Sample characteristic and mean values of chronological age and DNAm age estimates of each clock are presented for each cohort after quality control. Δage is defined by subtracting chronological age from DNAm age. Standard deviations are in parentheses unless otherwise defined. NLD = the Netherlands, SCT = Scotland, SWD = Sweden, UK = United Kingdom.

| Dataset | PI/Contact | Ancestry | Platform | Data type used | Total | Cases | Controls | Age (sd) |
|---|---|---|---|---|---|---|---|---|
| GSE4103 | RA Ophoff | Dutch | 27K | IDAT files | 624 | 337 | 287 | 33.3 (12.1) |
| GSE4119 | RA Ophoff | Dutch | 450K | IDAT files | 96 | 62 | 34 | 31.1 (10.2) |
| TBD | RA Ophoff | Dutch | 450K | IDAT files | 324 | 160 | 164 | 34.4 (11.4) |
| TBD | RA Ophoff | Dutch | 450K | IDAT files | 72 | 36 | 36 | 59.2 (5.7) |
| TBD | PF Sullivan | Swedish | 450K | IDAT files | 96 | 48 | 48 | 59.8 (5.9) |
| GSE80417 | A McQuillin | Scottish | 450K | (un)methylated intensities | 847 | 414 | 433 | 44.6 (12.9) |
| GSE84727 | D St. Clair | UK | 450K | (un)methylated intensities | 675 | 353 | 322 | 40.4 (15.0) |
| Total | | EUR | 27/450K | Mixed | 2,734 | 1,410 | 1,324 | 40.4 (28.1) |

**Table S2. Overview of datasets included in study.** Multiple datasets of whole blood DNAm data across four European cohorts were included in the study. Shown above are some sample characteristics and accompanying GEO accession numbers for each dataset before quality control. PI = Principal Investigator, UK = United Kingdom, EUR = European.

| Dataset | Ancestry | Platform | Tissue | Data type | Total | Cases | Controls | Age (sd) |
|---|---|---|---|---|---|---|---|---|
| GSE74193 | AA/EUR | 450K | DLPFC | IDAT files | 503 | 224 | 279 | 46.9 (15.4) |
| GSE61107 | - | 450K | Frontal cortex | IDAT files | 48 | 24 | 24 | 61.7 (19.2) |
| GSE61380 | - | 450K | Frontal cortex | (un)methylated intensities | 33 | 18 | 15 | 44.0 (15.7) |
| GSE61431 | - | 450K | Frontal cortex | (un)methylated intensities | 43 | 20 | 23 | 61.8 (17.5) |
| Total | Mixed / unknown | 450K | Frontal cortex | Mixed | 627 | 286 | 341 | |

**Table S3. Overview of datasets included in brain analysis.** Multiple datasets of postmortem brain DNAm data were included in in the analysis. Shown above are some sample characteristics and accompanying GEO accession numbers for each dataset before quality control. AA = African American, EUR = European, DLPFC = Dorsolateral prefrontal cortex.

**lm(formula = Δage ~ Dataset + Ethnicity + Platform + Age.continuous + Status*Age.groups*Sex)**

| | Horvath Δage | | | | Levine Δage | | | |
|---|---|---|---|---|---|---|---|---|
| **Model variables** | **Df** | **Sum Sq** | **Mean Sq** | **P-value** | **Df** | **Sum Sq** | **Mean Sq** | **P-value** |
| Dataset | 6 | 1795 | 299.2 | 2.0E-15 | 6 | 679 | 113.2 | 6.7E-03 |
| Ethnicity | - | - | - | - | - | - | - | - |
| Platform | - | - | - | - | - | - | - | - |
| Age.continuous | 1 | 116 | 115.8 | 2.1E-02 | 1 | 1054 | 27.7 | 1.5E-07 |
| Sex | 1 | 46 | 45.7 | 1.5E-01 | 1 | 412 | 412.2 | 1.0E-03 |
| Age.group | 4 | 645 | 161.3 | 6.1E-06 | 4 | 512 | 128.0 | 9.3E-03 |
| Status | 1 | 288 | 288.0 | 2.8E-04 | 1 | 916 | 915.7 | 9.8E-07 |
| Age.group:Status | 4 | 227 | 56.7 | 3.4E-02 | 4 | 491 | 122.7 | 1.2E-02 |
| Status:Sex | 1 | 12 | 11.9 | 4.6E-01 | 1 | 51 | 50.9 | 2.5E-01 |
| Age.group:Sex | 4 | 329 | 82.1 | 4.5E-03 | 4 | 694 | 173.5 | 1.1E-03 |
| Age.group:Sex:Status | 4 | 66 | 16.3 | 5.5E-01 | 4 | 273 | 68.3 | 1.3E-01 |
| Residuals | 2135 | 46331 | 21.7 | - | 2137 | 81187 | 38.0 | - |

**Table S9. Results three-way interaction model of age, sex, and status on Δage.** Shown are the contributions of each variable in the three-way interaction model presented by an analysis of variance table. The full model is displayed in the top row. Age.groups are defined by decades. Ethnicity and Platform are collinear with Dataset and thus do not have output. Df = degrees of freedom; Sum Sq; sum of squares; Mean Sq; mean of squares; P-value corresponds to the F-test in the anova() function.

| Sex | Age interval | Controls | Cases | Clock | Direction | β (Δage) | 95% CI | P |
|---|---|---|---|---|---|---|---|---|
| **Women** | **<25** | 84 | 28 | Horvath | decelerated | -2.36 | -4.07 — -0.64 | 7.3E-03 |
| **Women** | **25-29** | 75 | 28 | Levine | accelerated | 3.12 | 0.67 — 5.64 | 1.3E-02 |
| **Women** | **>36** | 217 | 192 | Levine | accelerated | 3.21 | 1.93 — 4.50 | 1.3E-06 |
| **Women** | **43-47** | 41 | 29 | Horvath | decelerated | -3.05 | -5.07 — -1.04 | 3.5E-03 |
| **Women** | **50-56** | 45 | 32 | Horvath | accelerated | 2.96 | 0.69 — 5.24 | 1.1E-02 |
| **Men** | **27-39** | 206 | 249 | Levine | accelerated | 1.72 | 0.62 — 2.81 | 2.3E-03 |
| **Men** | **<29** | 187 | 223 | Horvath | decelerated | -1.39 | -1.11 — 1.37 | 3.6E-03 |
| **Men** | **44-48** | 62 | 45 | Horvath | decelerated | -2.10 | -4.03 — -0.17 | 3.3E-02 |
| **Men** | **49-59** | 126 | 110 | Levine | accelerated | 2.44 | 0.67 — 4.21 | 7.1E-03 |

**Table S10. Sex-specific DNAm aging in schizophrenia is variable across chronological age.** For each identified change point and corresponding age interval, the number of samples and estimated Δage difference between cases and controls (β) with corresponding 95% confidence intervals (CI), and p-value (P) are presented. The sex, type of clock, and direction of aging effect are shown as well.

| | | Horvath Δage | | Levine Δage | |
|---|---|---|---|---|---|
| **Model variables** | **Model comparison** | **Δage $R^2$** | **P-value** | **Δage $R^2$** | **P-value** |
| Model x: baseline | | 3.9% | - | 3.1% | - |
| Model y: baseline (+ smoking) | - | 4.9% | - | 5.3% | - |
| Model z: baseline (+ cell types) | - | 8.2% | - | 22.1% | - |
| Model 0: baseline (+ smoking/cell) | - | 9.4% | - | 22.8% | - |
| Model 1: + status | Model 0 vs 1 | 9.3% | 0.86 | 22.8% | 0.26 |
| Model 2: + status*age.continuous | Model 1 vs 2 | 9.4% | 0.10 | 23.2% | 1.3E-03 |
| Model 3: + status*age.groups | Model 1 vs 3 | 10.6% | 2.1E-04 | 23.2% | 0.05 |
| Model 4: + status*age.groups*sex | Model 3 vs 4 | 10.8% | 0.15 | 23.6% | 0.05 |

**Table S11. Age- and sex-specific effects of DNAm aging in schizophrenia adjusted for smoking and cell type estimastes.** Shown are the contributions of interaction effects between disease status and age and sex on Δage when adjusted for DNAm smoking scores (baseline model y) and blood cell type proportions (baseline model z). The full baseline model is defined as Δage ~ dataset + ethnicity + age.continuous + sex + DNAm smoking score + DNAm blood cell type proportions. For other models, the variable(s) in addition to the full baseline variables are shown with the corresponding variance explained ($R^2$) in Δage. Interaction terms with chronological age are modeled as a continuous variable (age.continuous) or a categorical variable (age.groups). The latter uses previously defined decades. Model comparison is performed to assess if the contribution of an interaction term is significant compared to a model without that term. The chi-square test is used to test two models with corresponding p-value presented. The results of these analysis are shown for both the Horvath and Levine clock. These analyses included only 450K samples for which smoking scores and cell type estimates can be computed (N=1,621, 867 controls and 754 cases).

| **Women: >36 years (case = 165, control = 178)** | **Variable R2** | **Variable P** | **Variable R2 adjusted** | **P adjusted** |
|---|---|---|---|---|
| Model - all selected variables | 23.6% | 2.2E-08 | - | - |
| **Levine Δage** | **7.7%** | **6.0E-06** | **2.8%** | **3.3E-03** |
| Batch/Ethnicity | 1.6% | 5.2E-01 | 2.9% | 1.1E-01 |
| Smoking | 9.3% | 6.7E-07 | 4.7% | 1.6E-04 |
| CD8.naive | 0.1% | 5.8E-01 | 1.0% | 7.4E-02 |
| CD4.naive | 2.6% | 9.5E-03 | 0.2% | 4.6E-01 |
| CD8T | 6.2% | 5.5E-05 | 0.3% | 3.7E-01 |
| NK | 6.2% | 5.1E-05 | 0.0% | 7.2E-01 |
| Granulocytes | 8.7% | 1.5E-06 | 0.7% | 1.3E-01 |

| **All samples: <29 years (case = 152, control = 146)** | **Variable R2** | **Variable P** | **Variable R2 adjusted** | **P adjusted** |
|---|---|---|---|---|
| Model - all selected variables | 49.8% | 3.7E-32 | - | - |
| **Horvath Δage** | **3.1%** | **2.1E-03** | **0.6%** | **1.4E-01** |
| Batch/Ethnicity | 7.4% | 3.6E-05 | 0.6% | 5.4E-01 |
| Age | 0.0% | 8.3E-01 | 0.2% | 3.9E-01 |
| Sex | 8.9% | 1.2E-07 | 1.0% | 2.7E-02 |
| Smoking | 28.8% | 3.3E-23 | 18.6% | 2.3E-19 |
| CD8T | 3.7% | 7.8E-04 | 0.0% | 9.7E-01 |
| Granulocytes | 3.4% | 1.3E-03 | 0.4% | 1.7E-01 |
| CD8pCD28nCD45RAn | 5.3% | 5.3E-05 | 1.0% | 3.1E-02 |
| PlasmaBlast | 0.2% | 4.3E-01 | 0.6% | 7.8E-02 |
| NK | 7.8% | 7.9E-07 | 0.4% | 1.5E-01 |
| Bcell | 3.1% | 1.9E-03 | 0.0% | 7.3E-01 |
| Mono | 2.8% | 3.2E-03 | 0.5% | 1.1E-01 |

**Table S12. DNAm aging significantly contributes to schizophrenia independent of smoking and cell types.** Shown are variables that significantly explain variance in SCZ disease status, selected by a penalized logistic regression analysis. The top and bottom table present results for women >36 years and all samples <29 years, respectively. Only samples assayed on the 450K platform were included as DNAm-based smoking scores and cell type proportions could be computed and included in the analysis. The top row of each table shows the proportion of variance explained in disease status (R^2) for all selected variables combined and the significance of a logistic regression model (glm, family="binomial") with each variable included compared to the null model of no variance explained. We also show the proportion of variance explained by each variable individually (Variable R2) and by each variable adjusted for all other selected variables (Variable R2 adjusted). The significance of Variable R2 adjusted is computed by comparing the model with all variables to a model with the variable of interest removed using the anova(test = "LRT") function. The result of this test is shown in the "P adjusted" column.

| All samples: <29 | Controls | Cases | Mean value in cases | β (Horvath Δage) | 95% CI | P |
|---|---|---|---|---|---|---|
| **Polygenic risk** | | | | | | |
| All - no stratification | 379 | 243 | 0.22 | -1.30 | -2.00 — -0.60 | 3.1E-04 |
| PRS1 - continuous | - | 243 | 0.22 | -0.35 | -0.77 — 0.07 | 1E-01 |
| PRS1 - low | 379 | 81 | -0.77 | -1.01 | -2.05 — 0.02 | 5.6E-02 |
| PRS1 - mid | 379 | 81 | 0.17 | -1.32 | -2.36 — -0.27 | 1.7E-02 |
| PRS1 - high | 379 | 81 | 1.25 | -1.58 | -2.62 — -0.54 | 3.0E-03 |

| | | | | | | |
|---|---|---|---|---|---|---|
| **Age of onset** | | | | | | |
| All - no stratification | 379 | 190 | 19.48 | -1.47 | -2.23 — -0.71 | 1.6E-04 |
| AOO - continuous | - | 190 | 19.48 | -0.03 | -0.21 — 0.15 | 7.3E-01 |
| AOO - early | 379 | 64 | 15.91 | -1.49 | -2.61 — -0.37 | 9.8E-03 |
| AOO - mid | 379 | 63 | 19.19 | -1.06 | -2.23 — 0.10 | 7.2E-02 |
| AOO - late | 379 | 63 | 23.41 | -1.85 | -3.03 — -0.68 | 2.0E-03 |

| | | | | | | |
|---|---|---|---|---|---|---|
| **Illness duration** | | | | | | |
| All - no stratification | 379 | 190 | 4.92 | -1.47 | -2.23 — -0.71 | 1.6E-04 |
| DUR - continuous | - | 190 | 4.92 | 0.06 | -0.14 — 0.25 | 5.7E-01 |
| DUR - short | 379 | 64 | 1.50 | -1.86 | -3.04 — -0.70 | 1.8E-03 |
| DUR - mid | 379 | 63 | 4.68 | -1.31 | -2.45 — -0.17 | 2.5E-02 |
| DUR - long | 379 | 63 | 8.63 | -1.23 | -2.38 — -0.08 | 3.6E-02 |

**Table S13. Integration of Horvath Δage with PRS, age of onset, and illness duration in early adulthood.** Analyses were performed by stratifying the analyses to only men and women <29 years of age. Only cases with available information were included in the analyses. Each phenotype was analyzed as both a continuous variable and as a categorical variable using equal tertiles from low to high bins. Mean values in cases for each phenotype are presented along with the association with Δage (β) and corresponding 95% confidence intervals and p-values. PRS1 = polygenic risk score PC1 (see Supplementary Information) scaled to mean zero with standard deviation of 1, AOO = age of onset, DUR = illness duration.
