## Supplementary Information for "Epigenetic age is accelerated in schizophrenia with age- and sex-specific effects and associated with polygenic disease risk"

**Table of contents**

**S1. Supplementary Materials and Methods**	**2**

S1.1 Cohort descriptions	2

S1.2 Data preprocessing and normalization	4

S1.3 DNA methylation clocks	5

S1.4 Statistical analyses	6

S1.5 Definition of age at onset across cohorts	7

S1.6 PRS1 rationale and explanation	8

S1.7 DNAm-based estimation of smoking	13

S1.8 DNAm-based estimation of blood cell types	13

**S2. Supplementary Results**	**14**

S2.1 Analysis of DNAm aging in relation to estimates of smoking and blood cell types	14

S2.2 Levine Δage affects schizophrenia independently from smoking and cell types in women	16

S2.3 Analysis of postmortem brain samples	17

**References**	**19**

### S1. Supplementary Materials and Methods

#### S1.1 Cohort descriptions

*The Netherlands*

The Dutch cohort is a case-control sample with inpatients and outpatients recruited from different psychiatric hospitals and institutions across the Netherlands, coordinated via academic hospitals in Amsterdam, Groningen, Maastricht, and Utrecht. Detailed medical and psychiatric histories were collected, including the Comprehensive Assessment of Symptoms and History (CASH), an instrument for assessing diagnosis and psychopathology. Only patients with a Diagnostic and Statistical Manual for Mental Disorders fourth edition (DSM-IV) diagnosis of schizophrenia were included as cases. All patients and controls were of Dutch descent, with at least three out of four grandparents of Dutch ancestry. The controls were volunteers and were free of any psychiatric history, the majority via the CASH. Whole blood DNA methylation data is collected for 586 cases and 516 controls. DNAm profiles were obtained using the Illumina’s Infinium 27k Human DNA methylation Beadchip (v1.2) and the Infinium 450k Human DNA methylation Beadchip (v1.2). The 27k dataset can be found on GEO (GSE41037) with more details elsewhere^1,2^. Part of the 450k samples can be found under GSE41169. We have furthermore assayed an additional 200 controls and 196 cases.

*Cohort description: University College London (UK)*

More details on the cohort can be found elsewhere ^3–5^. The UCL cohort is a case-control sample recruited from London and South England consisting of unrelated cases and ancestrally matched controls. All subjects were included if both parents were of English, Irish, Welsh or Scottish descent, with at least three out of four grandparents having the same origins. All cases were selected for having prior International Classification of Diseases 10 (ICD10) diagnosis of schizophrenia made by National Health Service (NHS) psychiatrists. The research subjects were then given interviews with the Schedule for Affective Disorders and Schizophrenia-Lifetime Version (SADS-L) schedule and further data were collected from NHS medical and nursing case notes and all other available sources. Therefore, all cases were selected on the basis of having a primary clinical diagnosis of schizophrenia made by a psychiatrist at interview according to ICD10 criteria and then at the probable level of schizophrenia with Research Diagnostic Criteria (RDC) made at interview by a second research psychiatrist. The control subjects were also interviewed with the initial clinical screening questions of the SADS-L and selected on the basis of not having a family history of schizophrenia, alcoholism or bipolar disorder and for having no past or present personal history of any RDC-defined mental disorder. Whole blood DNA methylation data (450K) is publicly available for 353 patients and 322 controls through GEO (GSE80417).

*Cohort description: Aberdeen (SCT)*

More details on the cohort can be found elsewhere^3–5^. The Aberdeen cohort is a case-control sample that contains patients diagnosed with schizophrenia and non-psychiatric controls who have self-identified as born in the British Isles (95% in Scotland). All cases met the DSM-IV and ICD-10 criteria for schizophrenia. Diagnosis was made by Operational Criteria Checklist (OPCRIT). Controls were volunteers recruited through general practices in Scotland. Practice lists were screened for potentially suitable volunteers by age and sex and by exclusion of individuals with major mental illness or use of neuroleptic medication. Volunteers who replied to a written invitation were interviewed using a short questionnaire to exclude major mental illness in the individual themselves and their first-degree relatives. Whole blood DNA methylation data (450K) is publicly available for 414 patients and 433 controls through GEO (GSE84727).

*Cohort description: Sweden (SWD)*

The Swedish cohort is a case-control sample that contains both patients and controls of older age, i.e. 50-70 years. Whole blood DNA methylation data (450K) is collected for 96 samples, for which after matching by predicted sex and genotype information, 37 cases and 32 controls were included in the analysis.

#### S1.2 Data preprocessing and normalization

IDAT files and thus raw fluorescence intensity values were available for the Dutch and Swedish cohorts while for the UK and Scottish cohorts methylated and unmethylated intensity values were downloaded from GEO data repository (Table S1). To analyze DNAm quantifications, we employed three data processing pipelines to accommodate the 450K and 27K arrays with raw data available and the 450K arrays with only methylation intensity values from GEO. For samples with IDAT files available, raw intensity values were read into an RGChannelSetExtended object in the R programming environment using the read.metharray() function in the minfi package (v1.20.2)^6^. For samples with IDAT files available (Dutch and Swedish cohort, n=1,212), we used probe detection P-values to exclude outlying samples. That is, samples with more than 5% of probes detected at P > 0.05 were excluded from further analyses (n=13). To capture technical variation, surrogate variables that cumulatively explained >90% of variation of the control probe intensity levels were estimated using the ctrlsva() function of the ENmix package (v1.10)^7^ and stored to account for technical variance in downstream analyses. For the 450K arrays, background distributions were estimated using 600 chip internal control probes and dye-bias correction applied using the “RELIC” procedure^8^ implemented through the preprocessENmix() function. Probe design type correction was applied by Regression on Correlated Probes (RCP) through the rcp() function of the ENmix framework^9^. For the 27K arrays, normal-exponential out-of-band (noob) correction, which applies background correction with dye-bias normalization, was applied using the preprocessNoob() function in minfi^10^. For 450K arrays with only methylated and unmethylated intensity values available, background distributions were estimated and adjusted separately for each color channel and probe type using preprocessENmix() and probe design type correction subsequently applied using RCP.

#### S1.3 DNA methylation clocks

DNA methylation (DNAm), the addition of a methyl group to a cytosine nucleotide primarily at cytosine-phosphate-guanine (CpG) sites in the genome, is variable across the lifespan and has been shown to change relatively consistently between individuals^1,11,12^. Variation across the epigenome can be aggregated and used to generate a highly accurate estimate of chronological age, known as DNAm age or the “epigenetic clock”^13–15^. The Hannum clock uses 71 probes with regression weights determined through a training dataset of whole blood 450K DNA methylation data in 656 adult samples, aged 19 to 101. The Hannum estimator accurately predicts age in whole blood samples of adults. The Horvath clock uses 353 CpGs, present on both the 27K and 450K array, with regression weights determined using 8,000 samples across 30 tissues and cell lines from children and adults across the lifespan, age 0 to 100 years (mean = 43, SD = 25). The Horvath estimator accurately predicts age across the lifespan and across tissues, including blood and brain. The Levine clock uses 513 CpGs, present on both the 27K and 450K array, with regression weights obtained by regressing the weighted average of ten routine clinical parameters, including age, on DNA methylation levels in whole blood of on average older adult samples. This generates an estimate of so called “phenotypic age” that is predictive of age in whole blood of older adults.

For each sample and each clock, DNAm age was estimated by incorporating DNAm levels of the pre-selected set of probes, identified by each estimator as predictive of chronological age, into a mathematical model that weighs each probe’s methylation value and sums it to an aggregate DNAm age estimate. The Hannum and Horvath clock estimated DNAm age on average closer to chronological age (mean Δage Hannum = 1.0, Horvath = 1.7), while the Levine clock (i.e. phenotypic age) underestimated age by 7.7 years. See Table S1 for more details. The Levine clock was trained in an older adult population on a surrogate measure of biological age, generated through a Cox regression optimized to identify mortality-associated variables^15^. It therefore underestimates chronological age at younger ages.

#### S1.4 Statistical analyses

For each clock and each cohort, we implemented a multivariable regression model predicting Δage as a function of schizophrenia status, sex, and age (model 1) and as well as predicting Δage as a function of sex and the interaction between schizophrenia status and chronological age (model 2). Chronological age and sex were included as covariates in all analyses, unless stated otherwise. Regression models were set up in R as follows;

model 1: lm(Δage ~ sex + age + **status**)

model 2: lm(Δage ~ sex + age + status + **status:age**)

For the Dutch cohort, batch and array platform were also included as covariates, as this cohort consists of multiple datasets from both the 27K and 450K platform. For each clock, regression coefficients with corresponding standard errors for each of the four cohorts were then supplied to the rma() function of the metafor package^16^ in R to fit a meta-analytic fixed-effect model with inverse-variance weights and obtain an overall effect size and test statistic.

To quantify the significance of age- and sex-specific effects, we determined the contribution of interaction effects on top of the main disease effect. We first combined all cohorts to maintain necessary sample sizes across age and sex groups. Age categories were defined by grouping samples by decades with ages 18 and 19 included in the first decade (18-30, 31-40, etc.). To quantify the gain in variance explained in Δage, models with the interaction term were compared to a baseline model without the interaction term.

model 3: lm(Δage ~ cohort + sex + age.groups + status + **age.group:status**)

model 4: lm(Δage ~ cohort + sex + age.groups + status + age.groups:status + sex:status + age.group:sex + **age.group:sex:status**)

#### S1.5 Definition of age at onset across cohorts

*The Netherlands*

Age at onset is calculated preferentially by first using the date of first reported psychotic episode. If this information is not available the following order of variables is used as substitute; date of start treatment due to psychosis, date of start treatment with antipsychotic medication, date of first psychotic problems determined through the Comprehensive Assessment of Symptoms and History (CASH). Age at onset is available for 148 patients with overlapping DNAm data.

*Cohort description: University College London (UK)*

Age at onset is defined by the Operational Criteria Checklist (OPCRIT) and available for 321 patients.

*Cohort description: Aberdeen (SCT)*

Age at onset is available for 241 patients.

#### S1.6 PRS1 rationale and explanation

Polygenic risk scores (PRS) were generated by extracting each SNP’s log odds ratio from an independent discovery GWAS of SCZ and applying these weights to SNPs in a second target dataset. For each individual, PRS is established by calculating the sum across weighted SNPs. As our cohorts were included in the discovery GWAS, the weights were derived from a leave-one-out analysis with that specific cohort excluded. Risk score profiles were calculated across ten GWAS P-value thresholds (5x10^-8^, 1x10^-6^, 1x10^-4^, 0.001, 0.01, 0.05, 0.1, 0.2, 0.5, and 1.0), as previously described^17^. While there is a rich correlation structure across p-value thresholds, the threshold with the most discriminatory power between cases and controls often differs between cohorts, which discourages from a one-size fits all approach. In addition, between-site variability and accompanying shifts in the distribution of polygenic risk can limit the interpretability of differential disease risk.

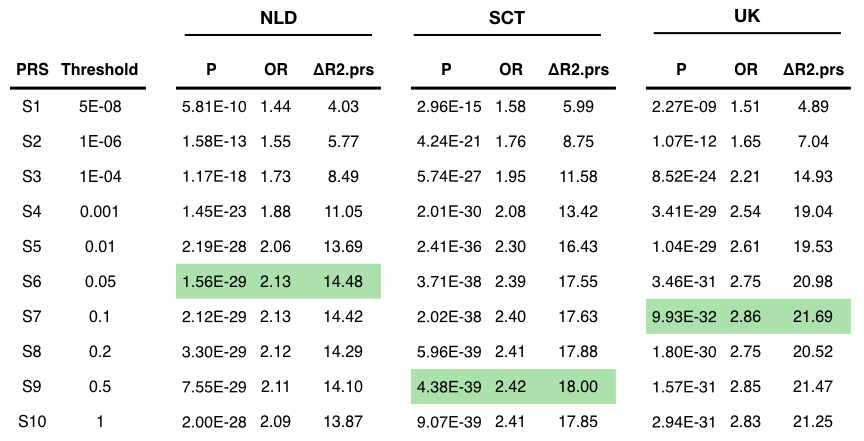

Each cohort with genetic data available has a different optimal GWAS p-value threshold that maximizes the magnitude of differential SCZ risk explained by PRS (highlighted in green). This demonstrates that choosing a single threshold across cohorts is suboptimal. NLD=Netherlands; SCT=Scotland, UK=United Kingdom, OR=odds ratio of PRS on disease status; ΔR2.prs=the unique variance explained in disease status by PRS.

Recent work using principal component analysis (PCA) in combination with normalization of the PRS across sites has shown to effectively eliminate between-site variation and concentrate disease risk information into the first principal component ^18^. This strategy works effectively because; (1) between-site variability can be eliminated by subtracting from all PRS values measured at one site the mean PRS among controls at that site, which aligns the PRS distribution with controls centered at zero; (2) PCA extracts disease relevant information across correlated thresholds of risk scores and concentrates this into a single component, which maximizes discriminatory power between cases and controls.

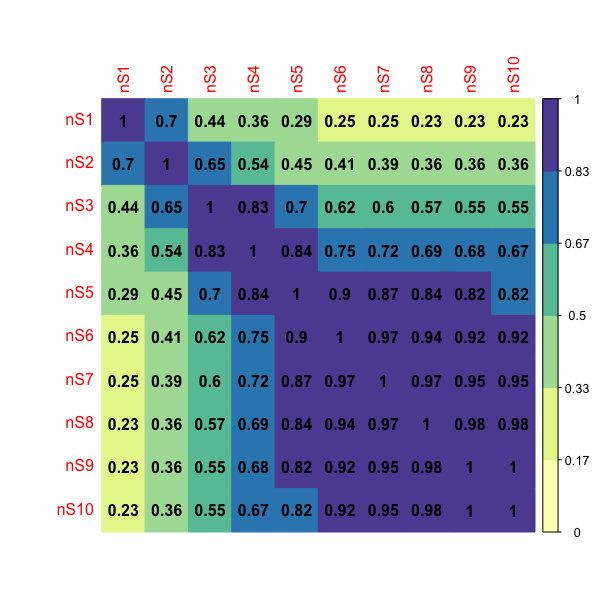

Correlation structure of SCZ PRS computed across various GWAS p-value thresholds (S1-10). While there is significant overlap between PRS of various thresholds, there is still unique information in each threshold that is not shared across threshold. The Pearson correlation is shown in each cell and color-coded as well. Principal component analysis can be used to extract disease risk information across thresholds and concentrating this in one dimension, representated by a single value.

We therefore adopted the analysis framework of Bergen et al, 2019 and first aligned the PRS distribution for each cohort and p-value threshold at mean zero for controls. We then performed PCA on all the normalized scores, i.e. across thresholds and cohorts, to concentrated disease risk in the first principal component and refer to this single value as “PRS1”. PRS1 explains 70.7% of the variance in PRS scores and captures 19.9% of the variance in disease status (OR=2.55) without adjusting for age and sex.

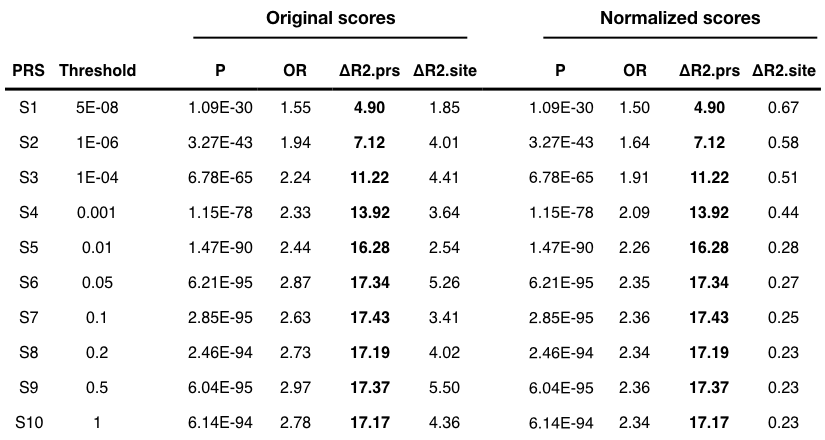

Between-site variability can be eliminated by subtracting from all PRS values measured at one site the mean PRS among controls at that site. Importantly, while accounting for cohort differences, this normalization does not affect the variance explained by PRS overall compared to the original scores (see ΔR2.site between original and normalized scores). OR=odds ratio of PRS on disease status; ΔR2.prs=the unique variance explained in disease status by PRS; ΔR2.site=the unique variance explained in disease status by cohort.

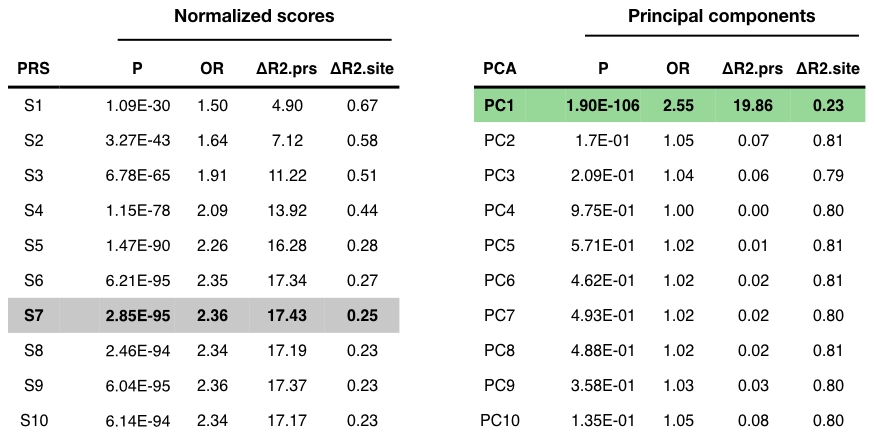

PCA on the normalized scores concentrates SCZ disease risk into the PC1 (highlighted in green). The discriminatory power by PC1 is superior to the original scores across various p-value thresholds. ΔR2.prs=the unique variance explained in disease status by PRS; ΔR2.site=the unique variance explained in disease status by cohort.

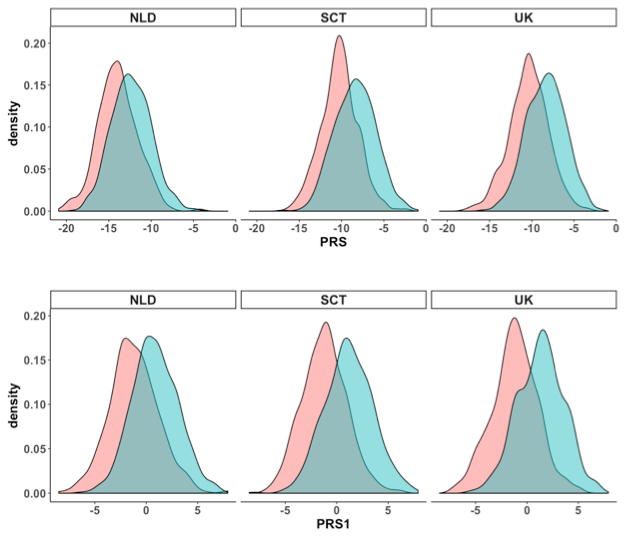

The distribution of PRS1 across cohorts (bottom panel) compared to PRS using the S6 P=0.05 threshold (top panel). PRS1 showns greater discriminatory power between cases and controls for each of the three cohorts.

#### S1.7 DNAm-based estimation of smoking

We computed a DNAm-based smoking score using CpG sites significantly associated with smoking. This smoking estimate has been shown to be a good proxy for smoking status^19^ and successfully used to account for the effects of smoking in large-scale DNAm studies of schizophrenia^5^. In a similar fashion as these studies, we calculate a weighted score across 183 DNA methylation sites, with the weights being effect sizes obtained from an epigenome-wide association study of smoking^20^.

#### S1.8 DNAm-based estimation of blood cell types

Estimated blood cell-type proportions were computed for all 450K array samples using the Horvath Methylation Age Calculator software. Further details on the derivation of estimates on CD8 T cells, CD4 T cells, nature killer cells, B cells, monocytes, and granulocytes can be found here^21^ and for plasma blasts, CD8+CD28-CD45RA- T cells, naive CD8 T cells here^13^.

### S2. Supplementary Results

#### S2.1 *Analysis of DNAm aging in relation to estimates of smoking and blood cell types*

Patients with SCZ smoke more than the general population^22^ and blood cell type composition changes across the lifespan^23^. To investigate the effect of these factors, we use DNAm-based smoking and cell type estimations (see Methods) as a proxy to evaluate their contribution to DNAm aging in SCZ.

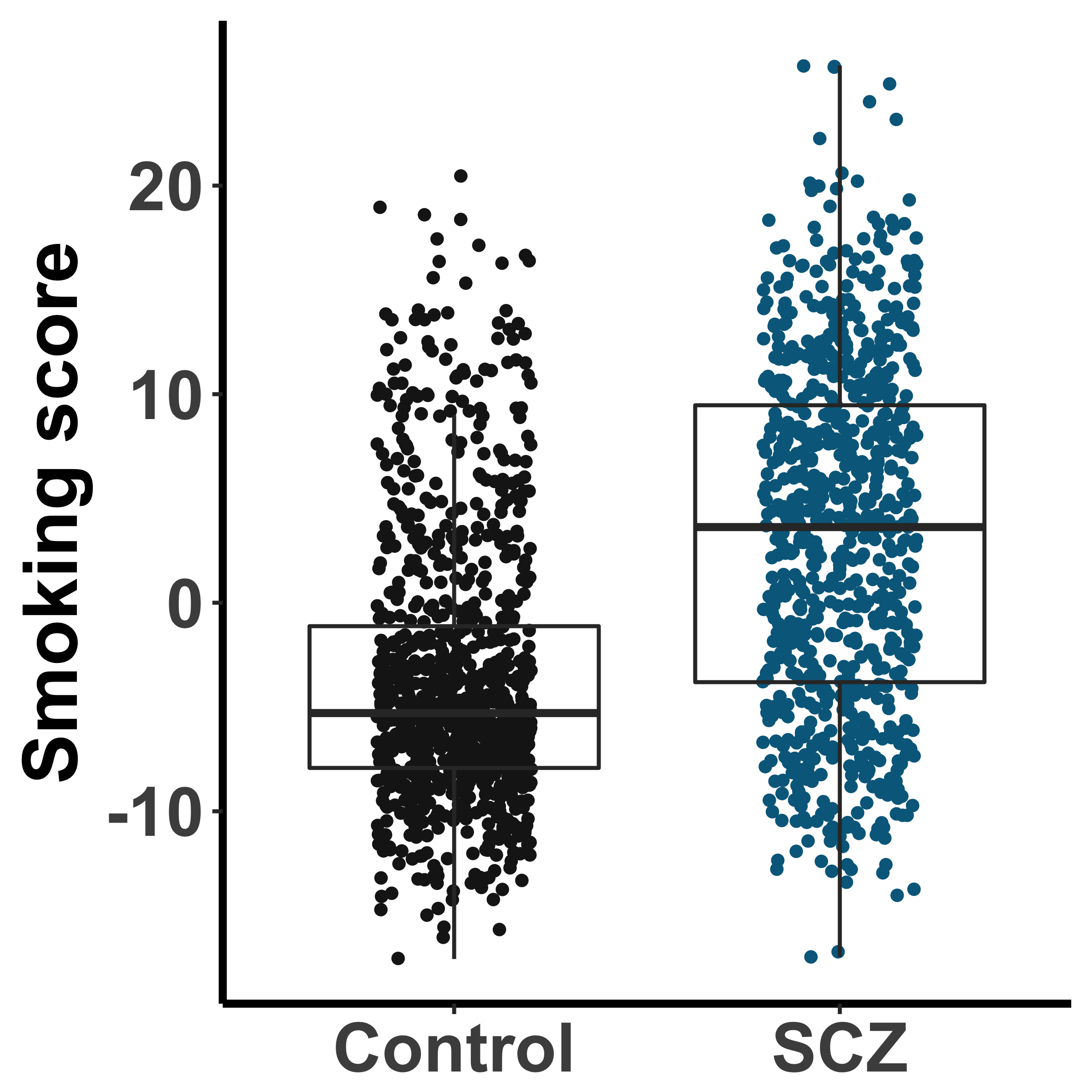

While DNAm clocks, by design, will encapsulate such effects, quantifying the contributions of each factor increases interpretability and helps understand the factors contributing to the differential aging findings. Here, we quantified the effects of DNAm smoking scores and blood cell types proportions on Horvath and Levine Δage in relation to the effect of disease status. For the Horvath clock, a baseline model with batch, ethnicity, sex, and age (as continuous) explains 3.9% of the variance in Δage. The addition of DNAm smoking scores and blood cell type proportions increases the fit of the model to 4.9% and 8.2%, respectively. The baseline model with both smoking scores and cell type estimates explain 9.4% of the variance in aging. While this reduces the main effect of disease status, the interaction between status and age (as categorical variable) remains significant and thus, as far as we can measure, independent of smoking and cell type composition.

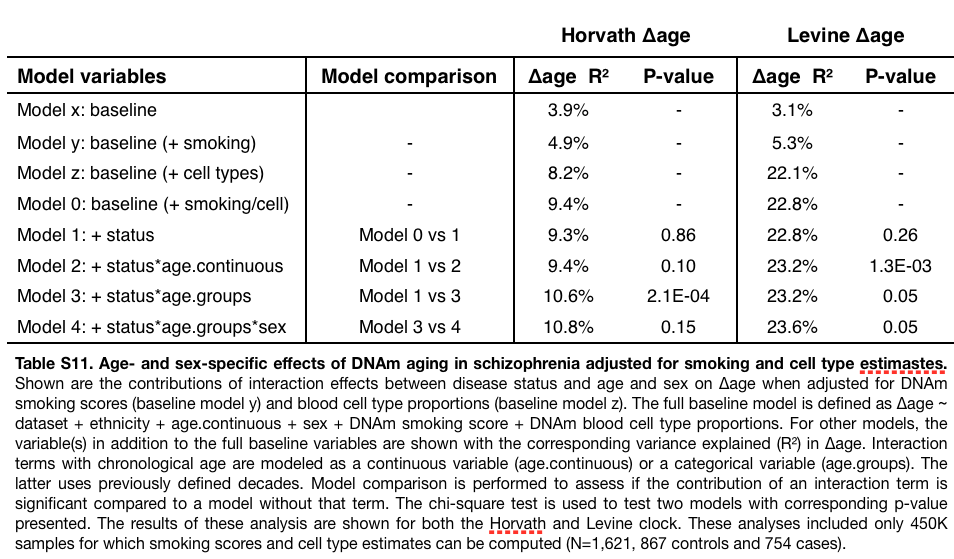

For the Levine clock, a baseline model with batch, ethnicity, sex, and age (as continuous) explains 3.1% of the variance in Δage. The addition of DNAm smoking scores and blood cell type proportions increases the fit of the model to 5.3% and 22.1%, respectively. The baseline model with both smoking scores and cell type estimates explain 22.8% of the variance in aging. This indicates that blood cell type composition explains a large proportion of the variance in Levine Δage, which is expected as the Levine clock is trained on blood mortality markers, including blood cell counts. Modeling smoking scores and blood cell type proportions as covariates reduces the main effect of disease status on Levine Δage. The interaction between status and age (as continuous variable) remains significant, similar to the analysis of Horvath Δage. For Levine Δage, the three-way interaction of status, age, and sex shows a slightly better model fit, after accounting for smoking and cell types, as well.

#### S2.2 Levine Δage affects schizophrenia independently from smoking and cell types in women

In women age 36 and older, the patient group in which we observed the most profound aging effects, a lasso logistic regression selected dataset/ethnicity, smoking, 5 cell types, and Levine Δage as independent variables to explain a total of 23.6% of the variance in SCZ disease status (P=2.2E-08) (Table S12). Levine Δage explains 7.7% individually and 2.8% (P=3.3E-03) when adjusted for other selected variables (Figure S10).

In individuals 29 years and younger, the group with significant Horvath deceleration aging, the lasso regression selected batch/ethnicity, age, sex, smoking, 7 cell types, and Horvath Δage as independent variabls to explain 49.8% of the variance in disease status (Table S12). A large proportion of this effect is driven by smoking, which explains 28.8%. Horvath Δage explains 3.1% of the variance in SCZ individually and 0.6% adjusted for other select variables (P=0.14). A significant proportion of the Horvath Δage effect on disease status is reduced by adjusting for smoking. However, smoking has no association with Horvath Δage in controls (Pearson r=0.01, P=0.95) nor in cases (Pearson r=-0.08, P=0.28) (Figure S11). As smoking covaries with SCZ disease status, it is difficult to distinguish these signals. In relation to SCZ genetic risk, smoking and blood cell types demonstrate limited effects on the observed pattern of differential aging across PRS1 (Figure S9).

#### S2.3 *Analysis of postmortem brain samples*

DNAm age estimates were generated for postmortem brain frontal cortex samples using publicly available data across four datasets (Table S2). The same data processing steps as described above were used for sample filtering and to generate beta values as input to estimate DNAm age. We excluded 16 samples due to missing data or discrepancy between reported and predicted sex and included only adult (age >= 18) frontal cortex samples, leaving 499 samples, 221 cases and 278 controls. Given the available sample sizes for some of the cohorts, we modeled Δage as a function of age and sex and disease status while correcting for cohort instead of performing a meta-analysis.

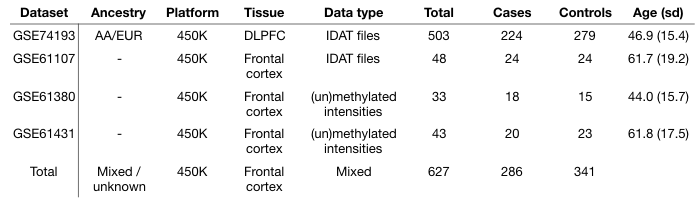

Multiple datasets of postmortem brain DNAm data were included in in the analysis. Shown above are some sample characteristics and accompanying GEO accession numbers for each dataset before quality control. AA = African American, EUR = European, DLPFC = Dorsolateral prefrontal cortex.

Only the Horvath clock yielded DNAm age estimates that closely correlated with chronological age. While the Hannum and Levine clock demonstrated decent correlations as well, they significantly underestimated chronological age. We therefore only investigated the Horvath clock, a multi-tissue estimator, and analyzed differential aging between cases and controls.

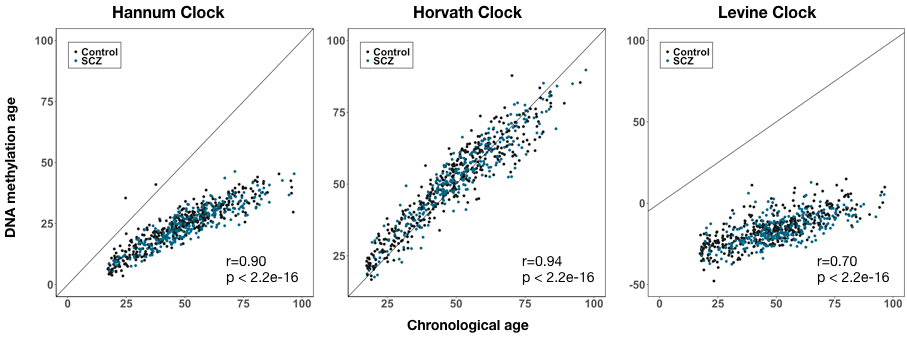
Correlation plots between DNAm age estimates and chronological age for postmortem brain samples. Results are shown for the Hannum (left), Horvath (middle), and Levine clock (right). Cases are colored blue and controls black. Pearson correlation and corresponding p-values are shown in the right bottom corner.

As with our analyses in blood, we first removed samples with descrepancies between reported sex and DNAm-based estimated sex (n=14) and samples with missing age information (n=2). Next, we regressed Δage on principal components of the control probes that explain >90% of the variation in control probe intensity values. The residuals were then added on mean(Δage) to generate Δage-adjusted, which preserves Δage in interpretable units (i.e. years). Using a multivariable linear regression model, we estimated differential DNAm aging in SCZ as follows;

model 1: lm(Δage ~ dataset + sex + age + **status**)

model 2: lm(Δage ~ dataset + sex + age + status + **status:age**)

Across the full dataset, we found no difference between cases and controls (ß=-0.29, P=0.46). We in addition did not observe a significant age-dependent disease effect, neither modeling age as a continuous variable (P=0.20) nor as a categorical variable (P=0.11). We also did not find Horvath age deceleration during early adulthood (age 18-30, ß=-0.09, P=0.92), like we observed in blood. We therefore conclude that there is no differential DNAm aging in the frontal cortex using postmortem brain samples.

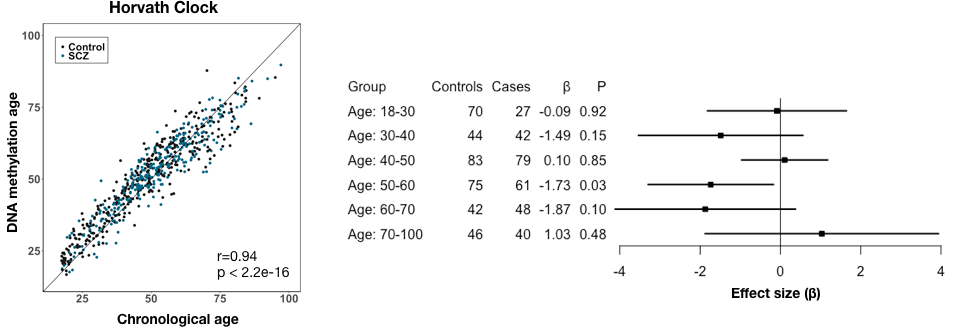

Results of Horvath DNAm aging in frontal cortex postmortem brain samples of SCZ cases and controls. The left plot shows the correlation between DNAm age estimates and chronological age. The right plot shows a forest plot of the aging effects (ß) between cases and controls across various age groups.
